# Inadvertent Transfer of Murine VL30 Retrotransposons to CAR-T Cells

**DOI:** 10.1101/2022.02.01.478686

**Authors:** Sung Hyun Lee, Yajing Hao, Tong Gui, Gianpietro Dotti, Barbara Savoldo, Fei Zou, Tal Kafri

## Abstract

For more than a decade genetically engineered autologous T-cells have been successfully employed as immunotherapy drugs for patients with incurable blood cancers. The active component in some of these game-changing medicines are autologous T-cells that express viral vector-delivered chimeric antigen receptors (CARs), which specifically target proteins that are preferentially expressed on cancer cells. Some of these therapeutic CAR expressing T-cells (CAR-Ts) are engineered via transduction with γ-retroviral vectors (γ-RVVs) produced in a stable producer cell line that was derived from murine PG13 packaging cells (ATCC CRL-10686). Earlier studies reported on the co-packaging of murine virus-like 30S RNA (VL30) genomes with γ-retroviral vectors generated in murine stable packaging cells. In an earlier study VL30 mRNA was found to enhance the metastatic potential of human melanoma cells. These findings raise biosafety concerns regarding the possibility that therapeutic CAR-Ts have been inadvertently contaminated with potentially oncogenic VL30 retrotransposons. In this study, we demonstrated the presence of infectious VL30 particles in PG13 cells conditioned media and observed the ability of these particles to deliver transcriptionally active VL30 genomes to human cells. Notably, VL30 genomes packaged by HIV-1-based vector particles transduced naïve human cells in culture. Furthermore, we detected transfer and expression of VL30 genomes in clinical-grade CAR-Ts generated by transduction with PG13 cells-derived γ-retroviral vectors. Our findings raise biosafety concerns regarding the use of murine packaging cell lines in ongoing clinical applications.

## Introduction

Following successful clinical trials the FDA and, later, the European, Canadian and Swiss regulatory administrations approved a gene therapy-based immunotherapy drug for adult patients with diffuse large B-cell lymphoma (DLBCL) who relapsed or did not respond to two conventional anticancer treatments. [1-3] The abovementioned medicine comprises autologous T-cells that were transduced *in vitro* with γ-RVVs expressing CARs directed to the B-cell specific protein CD19.

Production of the abovementioned therapeutic viral vectors is premised on a stable producer cell line derived from the murine PG13 packaging cells (ATCC CRL-10686). [4-6]. Reproducibility in the quality and quantity of vector preparations and the ability to scale up γ-RVV production were the impetus for the development of various stable packaging cell lines, most of which were derived from NIH 3T3 fibroblast cells. [7] Active murine endogenous retroviruses (ERVs) [8-10] raise biosafety concerns associated with the possibility that murine LTR-retrotransposons may be co-packaged along with clinical-grade γ-RVVs. [11] Indeed, earlier studies reported on efficient packaging of the murine VL30 retrotransposon into γ-RVV particles generated in various murine packaging cells. [12-17] Furthermore, a study by Song *et al*. demonstrated the ability of VL30 genomes to enhance the metastatic potential of human melanoma cells in immunodeficient mice. [18] In an earlier study, Purcell *et al*. detected VL30 genomes in lymphoma cells in non-human primates transplanted with γ-RVV-transduced hematopoietic stem cells. [16]

VL30 genomes do not include protein-encoding open reading frames. [18, 19] However, the VL30 mRNA functions as a long non-coding RNA (lncRNA), which efficiently binds the murine and the human tumor suppressor protein PTB-associated splicing factor (PFS). [18, 20-22] This protein is involved in multiple cellular pathways including DNA repair, RNA processing and regulation of the innate immune response. [23-27] We consider the possibility that inadvertent infection of human cells with VL30 retrotransposons can potentially mediate insertional mutagenesis, induce oncogenic pathways, and contribute to the emergence of novel pathogens. [28] Importantly, to date, the PG13 packaging cell line has not been characterized for retroelements secretion, and the FDA-required quality-control analysis of CAR-Ts does not include testing for the presence of endogenous retroelement genomes at the various stages of CAR-T production. In this study, for the first time, we characterized the secretion of VL30 genome-containing γ-RVV particles by PG13 packaging cells. Our findings indicate that murine VL30 genomes were efficiently delivered, reverse-transcribed and expressed in human 293T cells following exposure to conditioned media from PG13 cells. VL30 genomes that integrated into the chromatin of human cells were mobilized and transferred by HIV-based vectors to naïve human cells in culture. Furthermore, we detected transcriptionally active VL30 genomes in primary human T-cells (from healthy donors) following treatment with clinical grade CAR-expressing γ-RVVs. Our findings raise biosafety concerns regarding ongoing and past usage of murine packaging cell lines in clinical gene therapy applications.

## Results

### Transfer of transcriptionally active VL30 genomes by γ-RVV particles secreted from the PG13 packaging cell line to human embryo kidney (HEK) 293T cells

To test the hypothesis that productive VL30 particles released by PG13 packaging cells can infect human cells *in vitro*, the packaging cell line PG13 [6] was purchased from the American Type Culture Collection (ATCC CRL-10686). Conditioned media from PG13 cells was employed on HEK 293T cells. Treated 293T cells were cultured for 2 weeks (4 passages), after which genomic DNA and total mRNA were analyzed to determine the presence of transcriptionally active integrated VL30 genomes. As shown in Figure 1, qPCR and qRT-PCR assays readily detected integrated VL30 genomes and mRNA in the abovementioned treated 293T cells. Specifically, up to 1.5 copies of the VL30 genome were detected per 100 293T cells.

**Figure 1.**
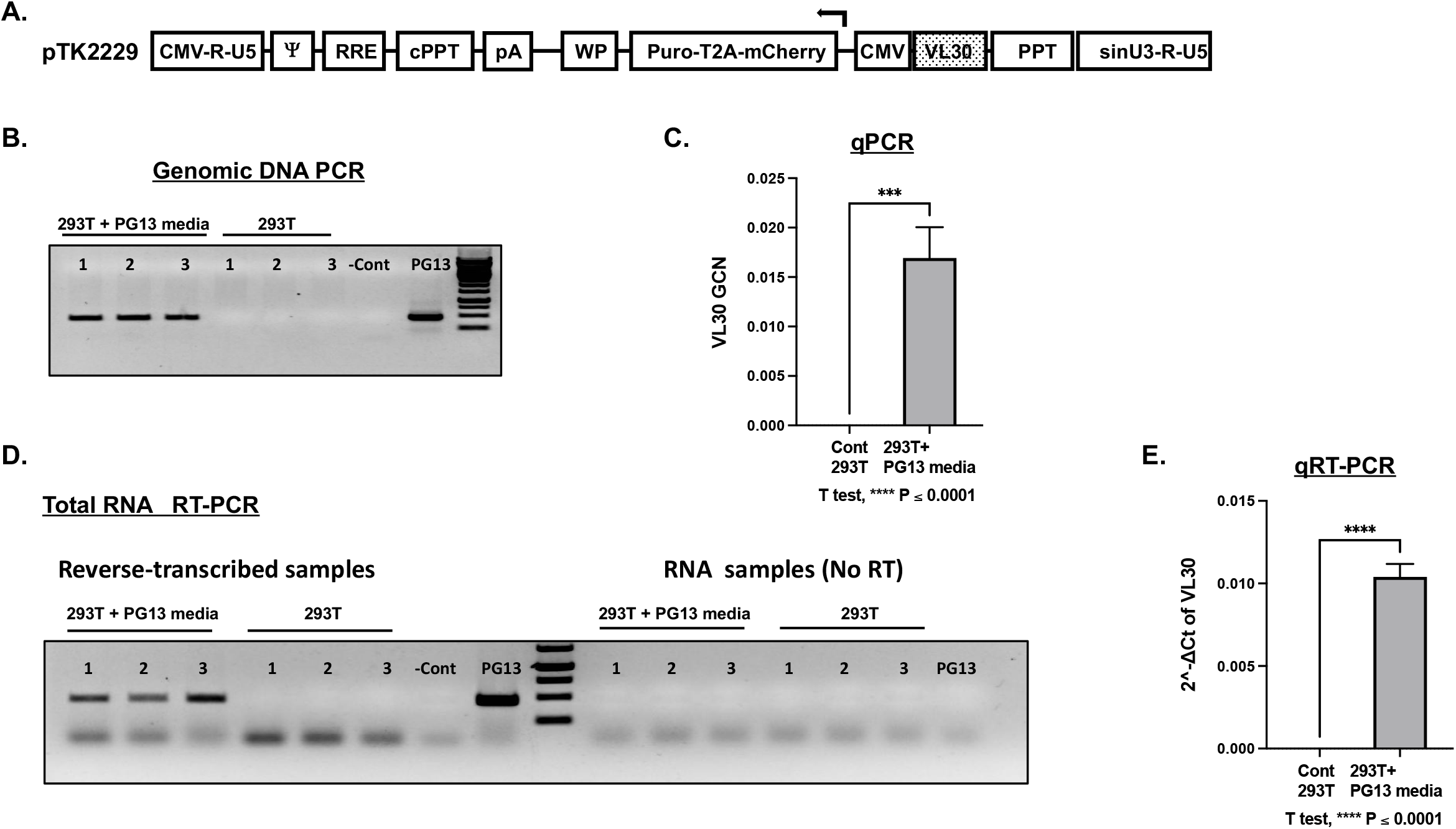
Inadvertently transferred VL30 genomes are transcriptionally active in HEK 293T cells Detection of VL30 genomes and mRNA in 293T cells following exposure to PG13 conditioned media. **A**. Physical map of the lentiviral vector cassette pTK2229 carrying a DNA sequence from the VL30 retrotransposon. The sequence is shown in a dotted box. It is located downstream of the internal CMV promoter, which is in opposite orientation to the LTRs and thus cannot initiate transcription of the VL30 sequence. The direction of transcription is indicated by an arrow. A CMV promoter replaces the 5’ U3. It is followed by the 5’ R and U5 regions. The packaging signal (**Ψ)**, Rev Response Element (RRE), central polypurine tract (cPPT), the woodchuck hepatitis virus post-transcriptional regulatory element (WP), the internal CMV promoter, and the Puro-T2A-mCherry reporter genes are shown. The modified self-inactivating (SIN) 3’ U3 is shown as SINU3. The parental 3’ poly purine tract (PPT) is shown. **B**. PCR-based analysis testing the presence of VL30 genomes in 293T cells. DNA was extracted from either naïve or treated (exposed to PG13 conditioned media) 293T cells following passages in culture. The presence of VL30 in the abovementioned DNA samples was determined by PCR. PCRs either in the absence of DNA or with DNA extracted from PG13 cells served as negative and positive controls. PCR products were visualized following gel electrophoresis. **C**. Bar graph showing vector copy number analysis of VL30 genomes in 293T cells following exposure to PG13 conditioned media. The gPCR -based assay was done in triplicate. Serial dilutions of DNA from pTK2229 vector transduced 293T cells served as a standard curve. DNA of naive 293T cells served as a negative control. **D**. RT**-**PCR-based analysis to characterize transcriptional activity of VL30 genomes in 293T cells. Total cellular RNA was extracted from either naïve or treated (exposed to PG13 conditioned media) 293T cells following passages in culture. The presence of VL30 mRNA in the abovementioned RNA samples was determined by RT-PCR. RT-PCR reactions in the absence of RNA or with RNA extracted from PG13 cells served as negative and positive controls. PCR of RNA samples prior to reverse-transcription served as controls for DNA contamination. PCR products were visualized following gel electrophoresis. **E**. Bar graph showing quantitative (q) RT-PCR analysis of VL30 mRNA in 293T cells following exposure to PG13 conditioned media. The gPCR based assay was done in triplicate. RNA of naive 293T cells served as a negative control. Levels of the endogenous hACTB mRNA served as an internal reference.

### PG13 cells’ conditioned media-mediated transfer of VL30 genomes to 293T cells is reverse transcription dependent

To support the hypothesis that the transfer of VL30 genomes to 293T cells following exposure to PG13 conditioned media is mediated by γ-RVVs, we tested the effects of reverse transcription inhibition on VL30 genome copy number in the abovementioned treated 293T cells. To this end PG13 cells were transduced with a γ-RVV (pTK2151, Fig 2-A) from which the green fluorescence protein (GFP) is expressed under the control of a simian virus 40 (SV-40) promoter. Conditioned media was obtained from naïve PG13 cells and from vector (pTK2151)-transduced PG13 cells and applied onto 293T cells, either in the presence or absence of 10µM azidothymidine (AZT), a nucleoside reverse transcriptase inhibitor. qPCR-based analysis VL30 genome copy number (GCN) in treated 293T cells demonstrated inhibition of VL30 transduction by AZT (Figure 2-B and Table 1). As expected, the presence of AZT efficiently inhibited the transduction of 293T cells by the pTK2151 γ-RVVs, which were generated either in the stable producer cell line PG13 or by transient three-plasmid transduction in 293T cells (Figure 2-C and Table 1).

**Table 1.** The effects of a reverse-transcriptase inhibitor of VL30 transduction. **A**. The table presents the raw data described in Figure 2-A. The conditioned media producing cell lines are outlined. The absence or presence of AZT (10µM) at the time of exposure to the abovementioned conditioned media is indicated by – and + signs, respectively. The copy number of VL30 genomes (GCN) in the abovementioned 293Ts is indicated. ND indicates VCN levels that were lower than the lower detection level by qPCR. The values of standard deviation (std) are shown. **B**. The table presents the raw data described in Figure 2-B. The conditioned media producing cell lines are outlined. The absence or presence of AZT (10µM) at the time of exposure to the abovementioned conditioned media is indicated by – and + signs, respectively. VL30 GCN in the abovementioned 293Ts is indicated. ND indicates VCN levels that were lower than the detection level obtained through qPCR. The values of standard deviation (std) are shown.

| A. |  |  |  | B. |  |  |  |
| --- | --- | --- | --- | --- | --- | --- | --- |
| Producing cell | AZT | VL30 GCN | std | Producing cell | AZT | γ-RVV VCN | std |
| PG13 | - | 0.0124 | 0.0011 | 293T +pTK2151 | - | 0.0682 | 0.0114 |
|  | + | ND |  |  | + | 0.0092 | 0.0016 |
| PG13 +pTK2151 | - | 0.0168 | 0.0068 | PG13 +pTK2151 | - | 0.0064 | 0.0002 |
|  | + | ND |  |  | + | ND |  |

**Figure 2.**
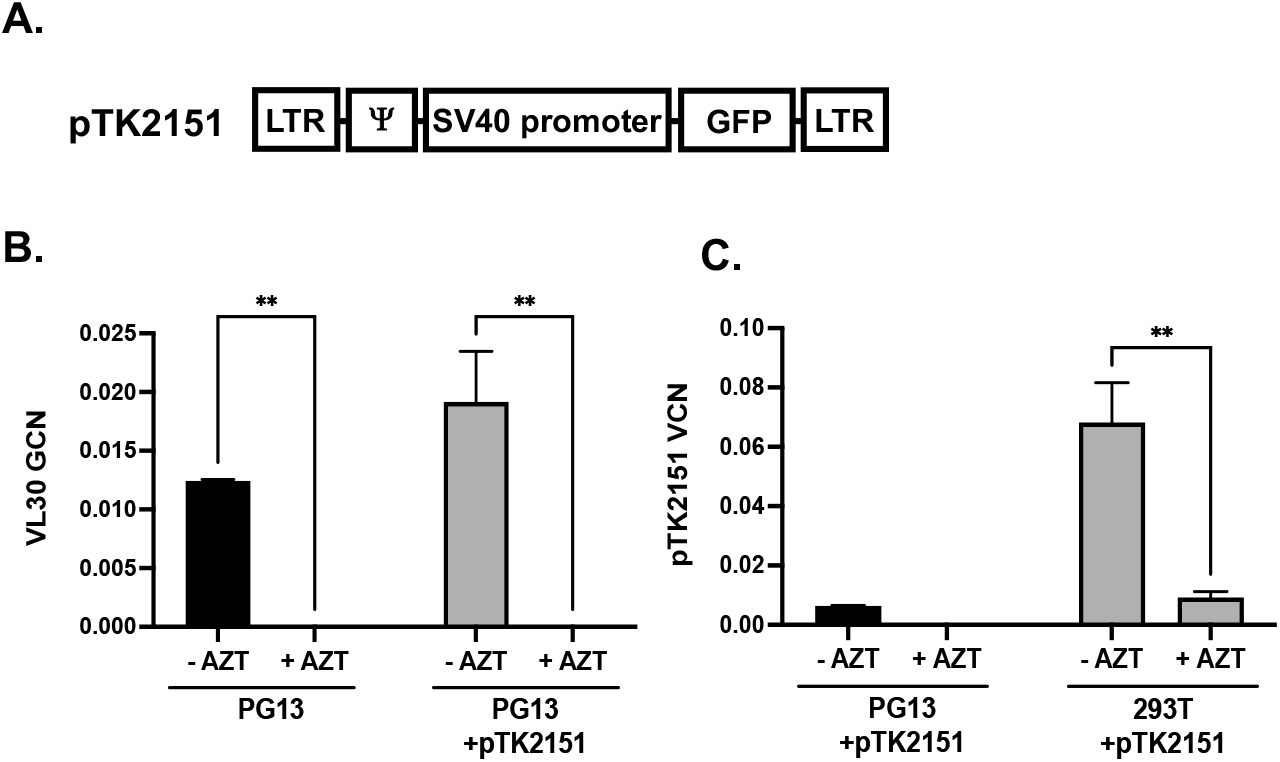
VL30 transduction of 293T cells is reverse transcription-dependent The effects of a reverse-transcriptase inhibitor on VL30 transduction. **A**. Physical map of the γ-retroviral vector (γ-RVV) pTK2151. The 5’ and 3’ non-SIN LTRs are shown. The packaging signal ()), the SV40 promoter and the GFP reporter gene are shown. **B**. Bar graph showing genome copy number (GCN) of VL30 in 293T cells following exposure to conditioned media collected from either PG13 or PG13 cells transduced with the γ-RVV pTK2151. Exposure to the abovementioned conditioned media was done either in the presence or absence of the reverse-transcriptase inhibitor azidothymidine (AZT 10 µM). The experiment was performed in triplicate. **C**. Bar graph showing VCN of the γ-RVV pTK2151 in 293T cells following exposure to conditioned media collected from either PG13 cells transduced with the γ-RVV pTK215, or 293T cells transiently transfected with the pTK2151 vector cassette, a VSV-g envelope and an γ-RVV packaging cassettes. Exposure to the abovementioned conditioned media was done either in the presence or absence of the reverse-transcriptase inhibitor AZT (10µM). The experiment was performed in triplicate Significance of the AZT effect on VL30 GCN and pTK2151 VCN was determined by 2-way ANOVA, *P≤0.05, **P≤0.01

### Increased VL30 genome copy number in 293T cells following co-culturing with PG13 cells

To better characterize VL30 genome function in human cells, we sought to increase VL30 genome copy number in 293T cells. To this end, we transduced 293T cells with the lentiviral vector pTK1261, which expresses the firefly luciferase and the GFP-blasticidin (BSD) selection marker under the control of the CMV promoter and the encephalomyelitis virus internal ribosome entry site (IRES), respectively. Vector-transduced cells were co-cultured with PG13 cells to confluency, at which point PG13 cells were eliminated by blasticidin selection and the vector transduced cells were re-cultured with fresh PG13 cells. The co-culturing/selection process was repeated 7 times, after which single-cell blasticidin-resistant 293T cell clones were isolated. DNA was isolated from a total of 8 single-cell clones and analyzed by qPCR for VL30 genome copy number. VL30 genomes were detected in 7 out of the 8 single cell clones. The range of VL30 GCN in the 7 VL30-positive clones was 2.40-5.17 (clones 8 and 6, respectively) VL30 genomes per 293T cell genome (Figure 3 and Table 2).

**Figure 3.**
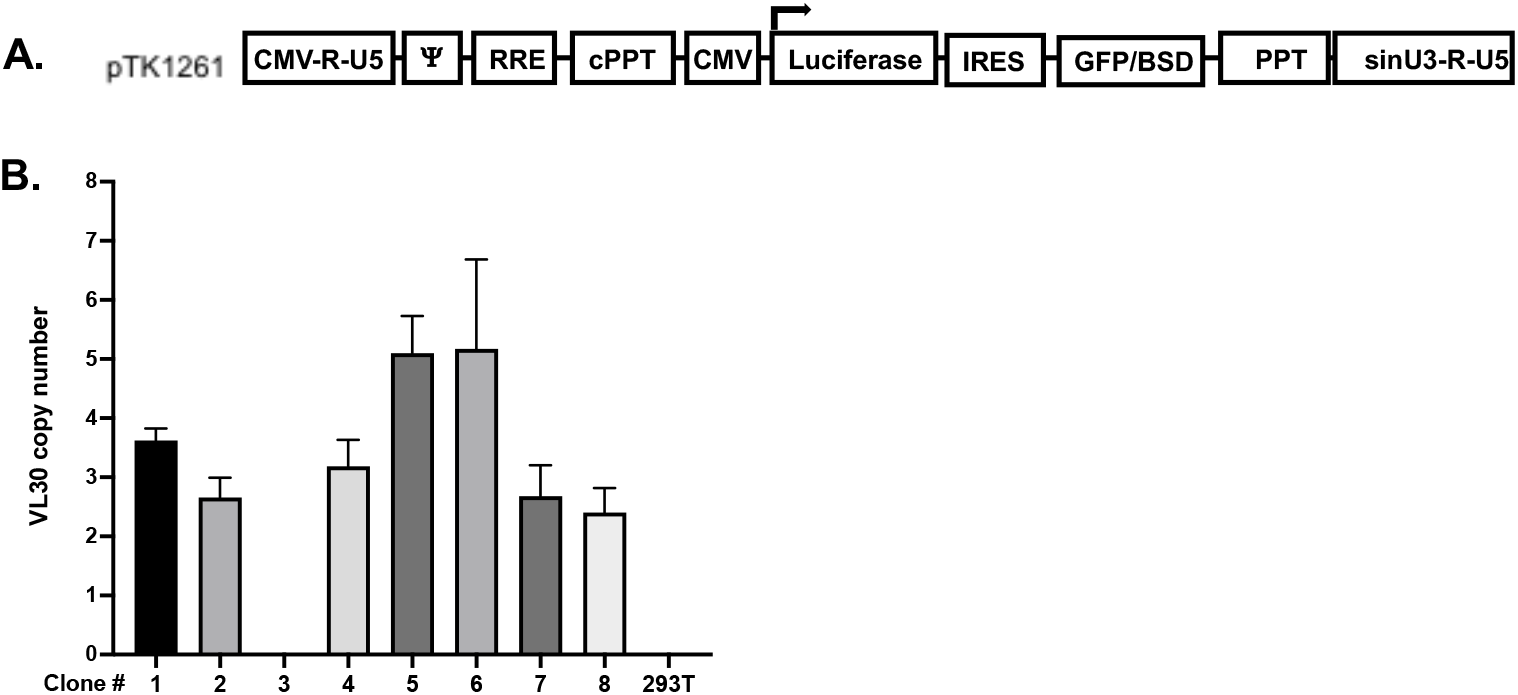
Copy number of VL30 genomes in single-cell clones of HEK 293T cells following co-culturing with PG13 cells. Increased VL30 genome copy number (GCN) in 293T cells, which were co-cultured with PG13 cells. **A**. Physical map of pTK1261 expressing the firefly luciferase and the GFP-blasticidin selection marker under transcriptional and translational control of a CMV promoter and an internal ribosome entry site (IRES) respectively. **B**. Bar graph showing VL30 GCN in single cell clones of 293T cells. PG13 and pTK2161 vector-transduced 293T cells were co-cultured for 7 passages. Single cell clones of co-cultured 293T cells were isolated in the presence of blasticidin (5 µg/ml). GCN in each cell clone was determined by qPCR. pTK2161-transduced 293T served as negative control. The experiment was performed in technical triplicate.

**Table 2.** Copy number of VL30 genomes in single-cell clones of HEK 293T cells following co-culturing with PG13 cells The table outlines the raw data of the experiments described in Figure 3-B. Clone numbers, values of VL30 GCN, and standard-deviations (std) are shown.

| Clone # | VL30 GCN | std |
| --- | --- | --- |
| 1 | 3.62 | 0.21 |
| 2 | 2.66 | 0.34 |
| 3 | ND |  |
| 4 | 3.18 | 0.45 |
| 5 | 5.10 | 0.63 |
| 6 | 5.17 | 1.51 |
| 7 | 2.68 | 0.52 |
| 8 | 2.40 | 0.42 |
| 293T | ND |  |

### Packaging of an endogenous murine retrotransposon into productive HIV-1 particles

To characterize the risk of VL30 genome dissemination within or between human subjects, we sought to evaluate the ability of HIV-1 vector particles to deliver VL30 genomes. To this end, two single cell clones of 293T cells (clones 6 and 7, which were isolated following the abovementioned co-culturing process with PG13 cells) were transiently transfected with a second-generation HIV-1 packaging cassette and a VSV-G envelope expression cassette. Conditioned media samples from the transiently transfected cells were employed on naïve 293T cells either in the presence or absence of 10µM AZT. At 24h after transduction, DNA samples were extracted and analyzed by qPCR to determine the presence of VL30 genomes. As shown in Figure 4 and Table 3, we readily detected VL30 genomes in 293T cells that were exposed to conditioned media of clones 6 and 7 in the absence of AZT. However, significant reduction in VL30 genome copy number was observed in 293T cells, that were exposed to the abovementioned conditioned media in the presence of AZT (an inhibitor of HIV-1 reverse transcriptase). These data suggest that HIV-1 particles can mediate the transfer of VL30 genomes between human cells. Furthermore, as shown in Tables 3 and 2, the level VL30 genome copy number in 293T cells treated with conditioned media from clones 6 and 7 (4.29 and 1.15 genome copies per 100 cells, respectively) correlated with the number of VL30 genomes in these cell clones (5.17 and 2.68 genomes per cell, respectively). To characterize the ability of HIV-1 reverse-transcribed VL30 genomes to integrate into human cells chromatin, we compared the VL30 genome copy numbers observed at 24 h (P0) and at 4 passages (P4) after transduction. As shown in Figure 5 and Table 4, in contrast to the γ-retroviral integrase, the HIV-1 integrase failed to integrate reverse-transcribed VL30 genomes into the chromatin of human 293T cells. This phenomenon could be explained by the lack of compatibility between the attachment sites (*att sequences*) at the VL30 LTRs and the HIV-1 integrase [29] and is in line with an earlier report describing the packaging of chimeric HIV-1/γ-RVV genomes by HIV-1 vector particles. [30]

**Figure 4.**
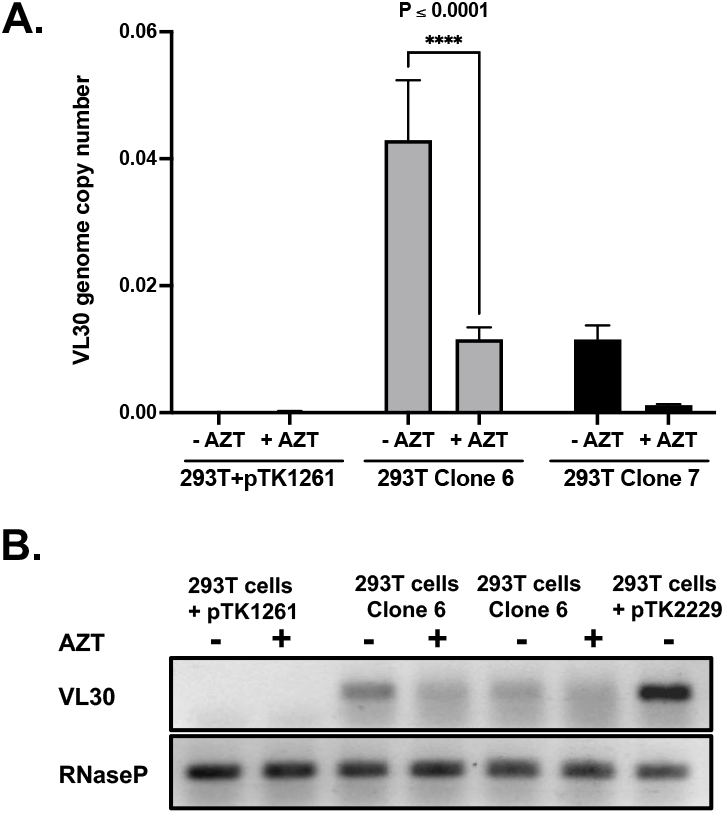
HIV-1 vector particles package and deliver VL30 genomes to naïve 293T cells VL30 transfer by human viral particles. **A**. To test the hypothesis that human pathogens, including HIV-1, can potentially transfer VL30 genomes, 293T cell clones 6 and 7 (VL30 genome copy number of 5.17 and 2.68, respectively) were transiently transfected with the HIV-1 vector packaging and the VSV-G envelope-expression cassettes. Vector particles were employed on naïve 293T cells either in the presence or absence of 10µM AZT. DNA was extracted from treated 293T cells and VL30 genome copy numb was determined by qPCR. **A**. Graph bar showing VL30 genome copy number in naive 293T cells exposed to lentiviral vector particles generated in cell clones 6 and 7, either in the presence or absence of 10 µM AZT. DNA samples extracted from 293T cells comprising the lentiviral vectors pTK1261 or pTK2229 served as negative and positive controls, respectively. Note that in the absence of AZT, the level of VL30 genome copy number in naïve 293T cells that were exposed to lentiviral vector particles generated in clone 6 and 7 (0.0429 ±0.0094 and 0.0115±0.0022, respectively) correlates with VL30 genome copy numbers in the respective vector producing single cell clones (Figure 3). The significant reduction in VL30 genome copy number in AZT treated 293T cells indicates that the observed transfer of VL30 genomes was reverse-transcription dependent. PCR amplification products of the endogenous RNaseP gene served as loading controls. P–values were determined by the 2-way ANOVA test. The experiment was performed in triplicate. **B**. Electrophoresis analysis of PCR products following amplification of DNA samples extracted from 293T cells transduced by HIV-1 particles generated in cell clones 6 and 7. DNA samples extracted from 293T cells transduced with the lentiviral vectors pTK1261 and pTK2229 served as negative and positive controls, respectively.

**Figure 5.**
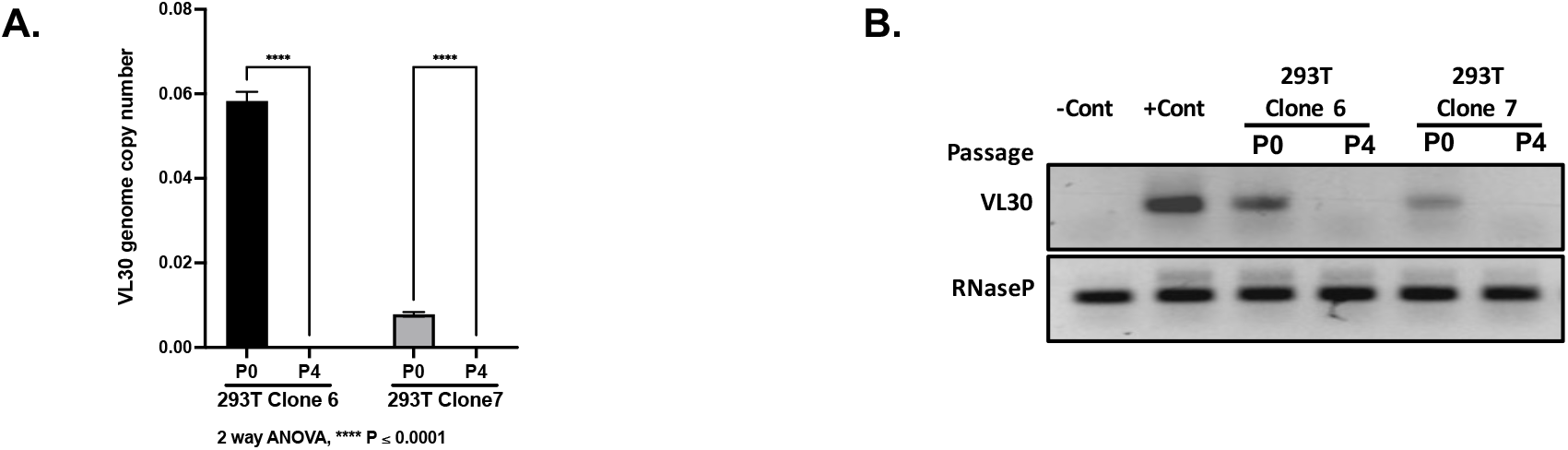
HIV-1 particles-delivered VL30 genomes are lost in passaged 293T cells. The effect of 293T cell replication on genome copy number of HIV-1 vector-transferred VL30. **A**. Bar graph showing VL30 genome copy number per cell in 293T cells treated with conditioned media containing HIV-1 vector particles packaged with VL30 genomes. Conditioned media samples were collected from cell clones 6 and 7 following transient transfection with a second-generation HIV-1 vector packaging plasmid, and a VSV-G envelope expression cassette. DNA samples of target 293T cells were analyzed by qPCR for the presence of VL30 genomes either at 24 h post-transduction (P0) or following 4 passages (P4). P-values were determined by the 2-way ANOVA test. The experiment was performed in triplicate. **B**. Electrophoresis analysis of PCR products following amplification of DNA samples extracted from 293T cells transduced by HIV-1 particles generated in cell clones 6 and 7. DNA samples extracted from 293T cells transduced with lentiviral vectors pTK1261 and pTK2229 served as negative and positive controls, respectively, and are shown as –Con and + Con, respectively. PCR amplification products of the endogenous RNaseP gene served as loading controls. P-values were determined by the 2-way ANOVA test. The experiment was performed in triplicate.

**Table 3.** Data of the qPCR assays described in Figure 4-A. The vector and the conditioned media employed on the target 293T cells are indicated. The presence or absence of 10 µM AZT at the time of transduction is indicated. DNA samples extracted from 293T cells transduced with the lentiviral vector pTK1261 served as negative controls. VL30 genome copy number per cell (GCN) in conditioned media-treated 293T cells and standard-deviations (std) are shown. PCR amplification products of the endogenous RNaseP gene served as loading controls. P –values were determined by the 2-way ANOVA test. The experiment was performed in triplicate.

| Cell line | AZT | VL30 GCN | std |
| --- | --- | --- | --- |
| 293T + pTK1261 | - | Not detected |  |
|  | + | Not detected |  |
| 293T + Clone 6 conditioned media | - | 0.0429 | 0.0094 |
|  | + | 0.0116 | 0.0018 |
| 293T + Clone 7 conditioned media | - | 0.0115 | 0.0022 |
|  | + | 0.0012 | 0.0002 |

**Table 4.**
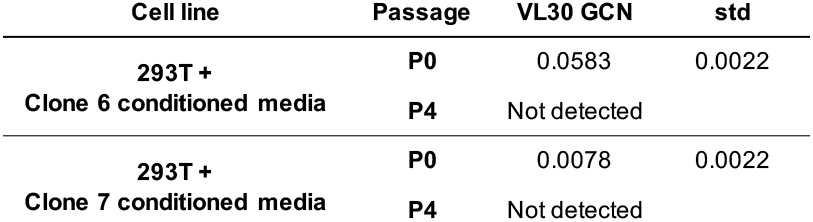
The table outlines the data employed in Figure 5-A. HIV-1 vector particles were generated in 293T cell clones 6 and 7 and employed on naive 293T cells. DNA was extracted from treated cells at either 24h (P0) or following 4 passages (P4) after treatment and analyzed by qPCR for VL30 genome copy number. The experiment was done in triplicate. The target cell line (293T cells) and the cell lines in which the HIV-1 particles containing conditioned media were generated are shown. The number of target cells passages (P0 and P4) are shown. Values of VL30 genome copy number per target cell and the standard deviation (std) are shown.

### Clinical grade γ–RVVs produced in the PG13 packaging cell line inadvertently transfer VL30 genomes to CAR-T cells

Next, we sought to evaluate the risk of inadvertent transfer of VL30 retrotransposons to human cells in a relevant clinical setting. To this end, we determine the presence of VL30 genomes in 8 populations of clinical-grade CAR-T cells and in 5 populations of naïve primary human T-cells. All cell populations were obtained from healthy human donors. CAR-T cells were generated by a clinical-grade transduction protocol using γ-RVV generated in PG13 producer cell lines, as described in [31]. A total of 13 samples of primary human cells were analyzed by qPCR for the presence of VL30 and γ–RVV genomes. DNA samples 3, 4, 8, 9 and 10 were extracted from control human T-cells that were not treated with the abovementioned γ-RVV (employed as negative controls). As shown in Figure 6 and Table 5, γ-RVV genomes were detected in all 8 populations of γ-RVV-treated T-cells. Vector copy number (VCN) of γ-RVV ranged from ∼13.5 to ∼0.6 vector copies per 100 T-cells (in samples 6 and 11, respectively). VL30 genomes were detected in 6 out of the 8 T-cell populations exposed to γ-RVV (cell populations 2, 5, 6, 7, 12 and 13). In these T-cell populations VL30 genome copy number (GCN) ranged from ∼40 to ∼0.5 genome copies per 1000 cells in samples 5 and 12, respectively. Two populations of γ-RVV-treated T-cells (samples 1 and 11) were observed to be negative for the presence of VL30 genomes. Importantly, there was correlation between γ-RVV VCN and VL30 GCN in the different T-cell populations (Figure 6E). This observation raises the possibility that low levels of VL30 genomes in samples 1 and 11, which exhibited the lowest levels of γ-RVV VCN, were below the qPCR detection level employed in this study.

**Table 5.**
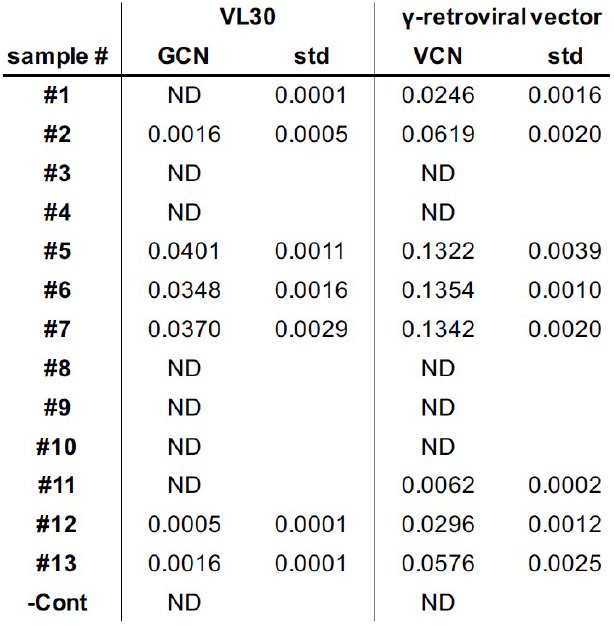
Genome copy number (GCN) and vector copy number (VCN) of VL30 and γ-RVV (respectively), in DNA samples extracted from primary human T-cells as determined by qPCR analysis. DNA samples 1, 2, 5, 6, 7, 11, 12 and 13 were extracted from human T-cells treated with the γ-RVV SFG.iC9.GD2.CAR.IL15. DNA of samples 3,4,8,9 and 10 were extracted from naïve primary human T-cells and served as biological negative controls. DNA extracted from naïve 293T cells served as technical negative control (-cont). ND indicates that the level of targeted sequences (VL30 and γ-RVV) in the tested DNA sample were under the level of detection.

**Figure 6.**
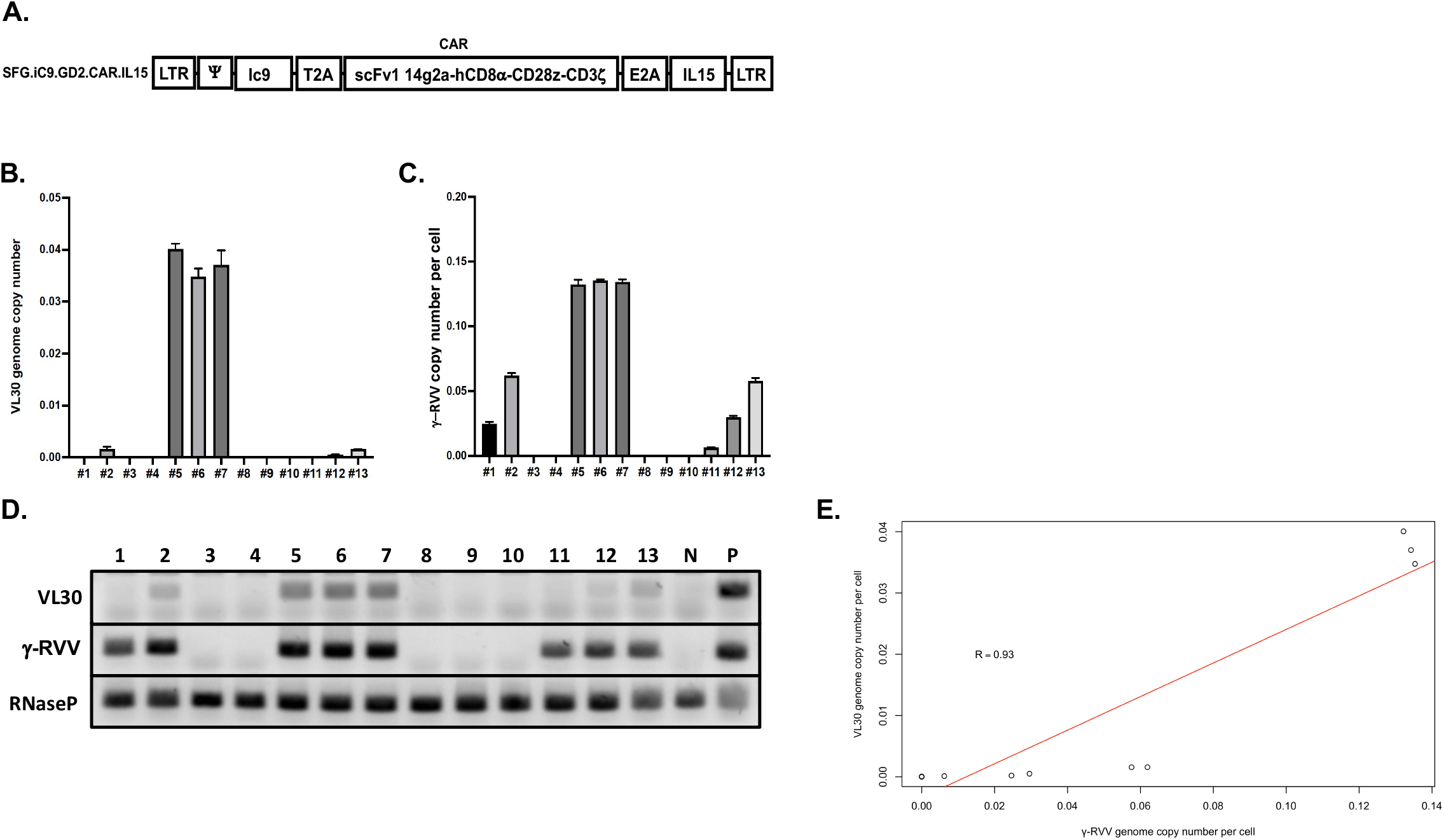
Transfer of VL30 genomes to primary human T-cells by γ-retroviral vector (γ-RVV) particles produced in PG13 cells qPCR-based analysis of VL30 genome copy number (GCN), and γ-RVV vector copy number (VCN) in primary human T-cells. Naïve primary human T-cells were treated with the chimeric antigen receptor (CAR)-carrying γ-RVV SFG.iC9.GD2.CAR.IL15 (samples 1, 2, 5, 6, 7, 11, 12 and 13). The vector was produced in the stable packaging cell line PG13. Samples 3,4,8,9 and 10 were not exposed to the abovementioned CAR-carrying γ-RVV and served as biologic negative controls. DNA samples extracted from the abovementioned control and vector-treated cells were used to determine GCN and VCN of the murine endogenous retrotransposon VL30 and the γ-RVV vector, respectively. DNA from naïve 293T cells served as a technical negative control. DNA from 293T cells transduced with the lentiviral vectors pTK2229 and pTK2151 served as positive control for VL30 and γ-RVV genome amplification, respectively. Amplification of the endogenous human gene hRNaseP served as loading control. The assay was performed in technical triplicate. **A**. A Physical map depicting the structure of the CAR–carrying γ-RVV, SFG.iC9.GD2.CAR.IL15. The vector’s non-self-inactivating (non-SIN) long terminal repeats (LTRs) and the packaging signal Ψ are shown. The inducible caspase-9 suicide gene, Ic9 is shown. The self-cleavable peptide from the thosea asigna virus (T2A) and the equine rhinitis virus (E2A) are shown. Sequences encoding the CAR including the single-chain variable fragment (scFv1) directed to the NB-antigen GD2 of the disialoganglioside GD2 (14g2a), the CD8α stalk and transmembrane domain, the CD28 intracellular domain and the CD3ζ chain are shown. The human IL15 cDNA containing sequence is shown. **B**. Graph bar showing GCN of VL30 in the abovementioned primary human T-cells. The experiment was performed in technical triplicate. **C**. Graph bar showing VCN of the γ-RVV SFG.iC9.GD2.CAR.IL15 in the abovementioned primary human T-cells. The experiment was performed in technical triplicate. **D**. Gel electrophoresis analysis of DNA amplification products of VL30, γ-RVV and hRNaseP sequences generated in the course of the above gPCR assay. DNA of naïve 293T cells (N) served as technical negative control for amplification of VL30 and γ-RVV sequences. DNA from 293T cells transduced with the lentiviral vectors pTK2229 and pTK2151 served as positive controls (P) for VL30 and γ-RVV genome amplification, respectively. **E**. Significant correlation between VL30 and γ-RVV genome copy number in human T-cells transduced with CAR-expressing γ-RVV. Linear regression graph demonstrating the relation between VL30 and γ-RVV genome copy number. The R value is indicated.

### VL30 genomes are transcriptionally active in primary human T-cells

Aware of the oncogenic potential of VL30 mRNA, we investigated the transcriptional activity of VL30 genomes in the abovementioned VL30-containing human T-cell populations. To this end, we employed the reverse-transcriptase (RT) qPCR assay on RNA samples extracted from T-cell populations 5, 6, and 7, as well as from populations 4 and 8, which were not treated with γ-RVVs and served as negative controls. As shown in Figure 7 and Table 6, VL30 transcripts were detected in all RNA samples from T-cell populations 5-7. These data suggest that VL30 genomes that were inadvertently transferred to primary human T-cells in the course of γ-RVV transduction were transcriptionally active.

**Table 6.** The table outlines the raw data described in figure 7. Values of 2^ ^ΔCt^ and standard deviation (std) are shown. ND indicate mRNA levels below the level of detection. P indicates positive control (RNA extracted from 293T cells transduced with the lentiviral vector pTK2229).

| sample # | $2^{-\Delta\Delta C_t}$ | std |
| --- | --- | --- |
| #4 | ND |  |
| #5 | 0.00053 | $4.98 \times 10^{-5}$ |
| #6 | 0.00048 | $3.10 \times 10^{-5}$ |
| #7 | 0.00052 | $1.34 \times 10^{-4}$ |
| #8 | ND |  |
| P | 0.03200 | $1.81 \times 10^{-3}$ |

**Figure 7.**
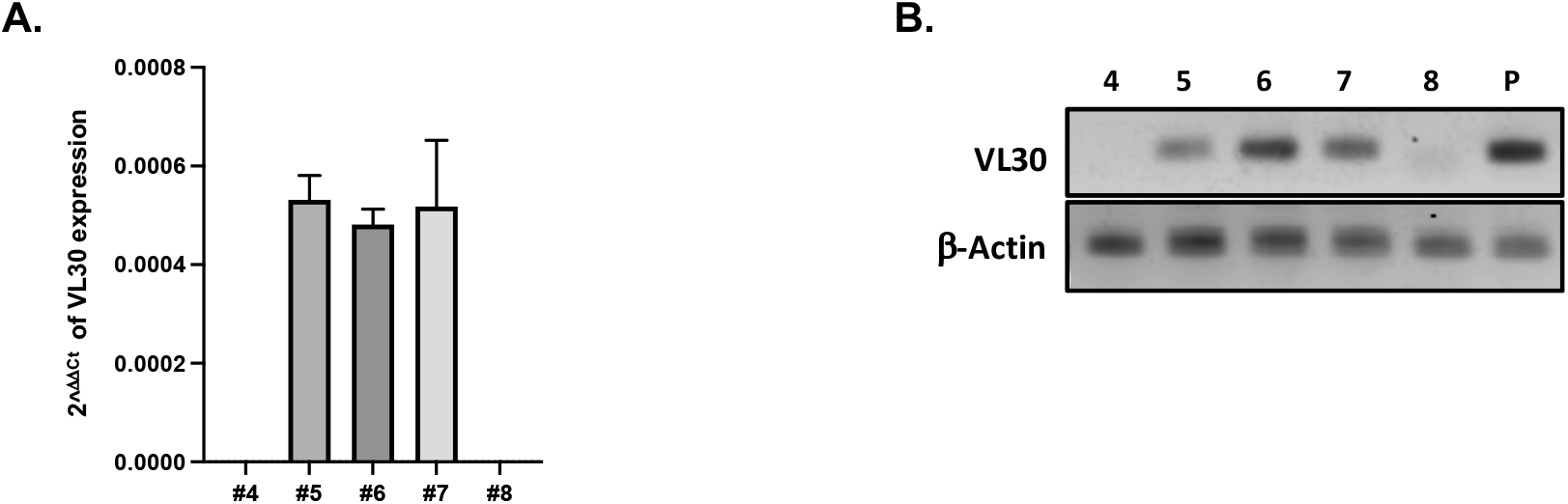
VL30 expression in human primary T-cells following transduction with γ-RVV vectors generated in PG13 cells. Quantification of VL30 expression in human T-cells. **A**. Bar graph showing VL30 expression relative to the endogenous β-actin gene. The qRT-PCR was performed in technical triplicate on RNA samples extracted from samples 4-8. Note that samples 4 and 8, in which VL30 genomes could not be detected (Figure 6), served as negative controls. **B**. Gel electrophoresis analysis showing the DNA products of the abovementioned qRT-analysis. Amplification products of RNA samples extracted from 293T cells transduced with the lentiviral vector pTK2229 (Figure 1) served as positive control (P). Amplification products on of the β-actin mRNA are shown.

### Short sequences of homology between the human genome and various VL30 genomes

The transcriptional activity of VL30 genomes in primary human T-cells and the ability of HIV-1 vector particles to mobilize VL30 genomes from human cells (Figure 4) increase the potential for recombination between VL30 genomes and either human or/and HIV-1 genomes. [28, 32-34] Prompted by this possibility, we searched for sequence homologies between various VL30 genomes and either the HIV-1 genome or the human genome. We could not detect sequence homology between VL30 genomes and the HIV-1 genome. Importantly, multiple short sequences of ∼30 bp in VL30 genomes were determined to be homologous to sequences throughout the human genome. Most of the abovementioned VL30-homologous human sequences were composed of interspersed repeats, including LTRs, LINEs, and SINEs, or simple repeats (Table 7).

**Table 7.**
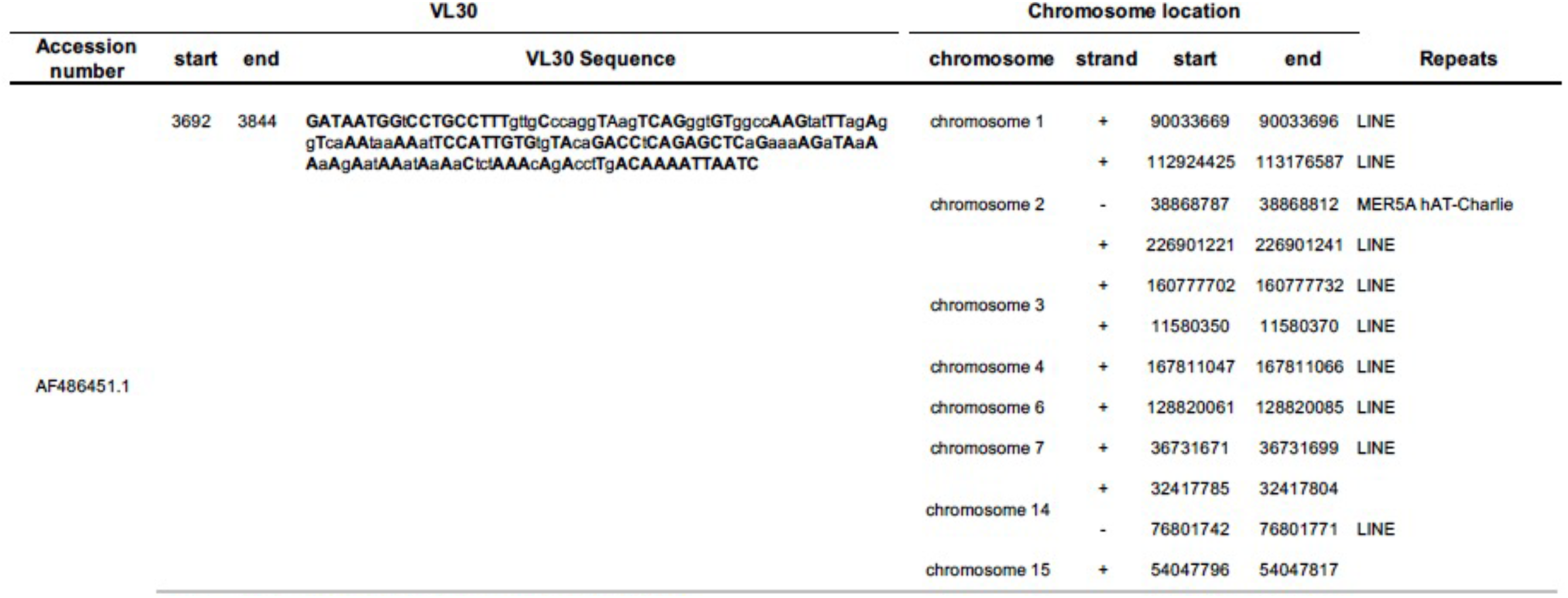
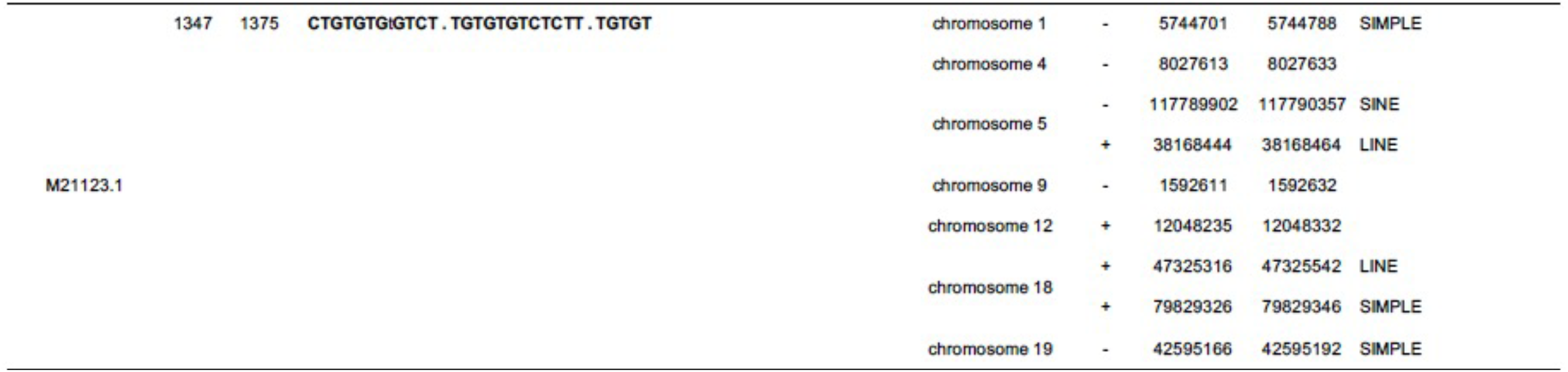
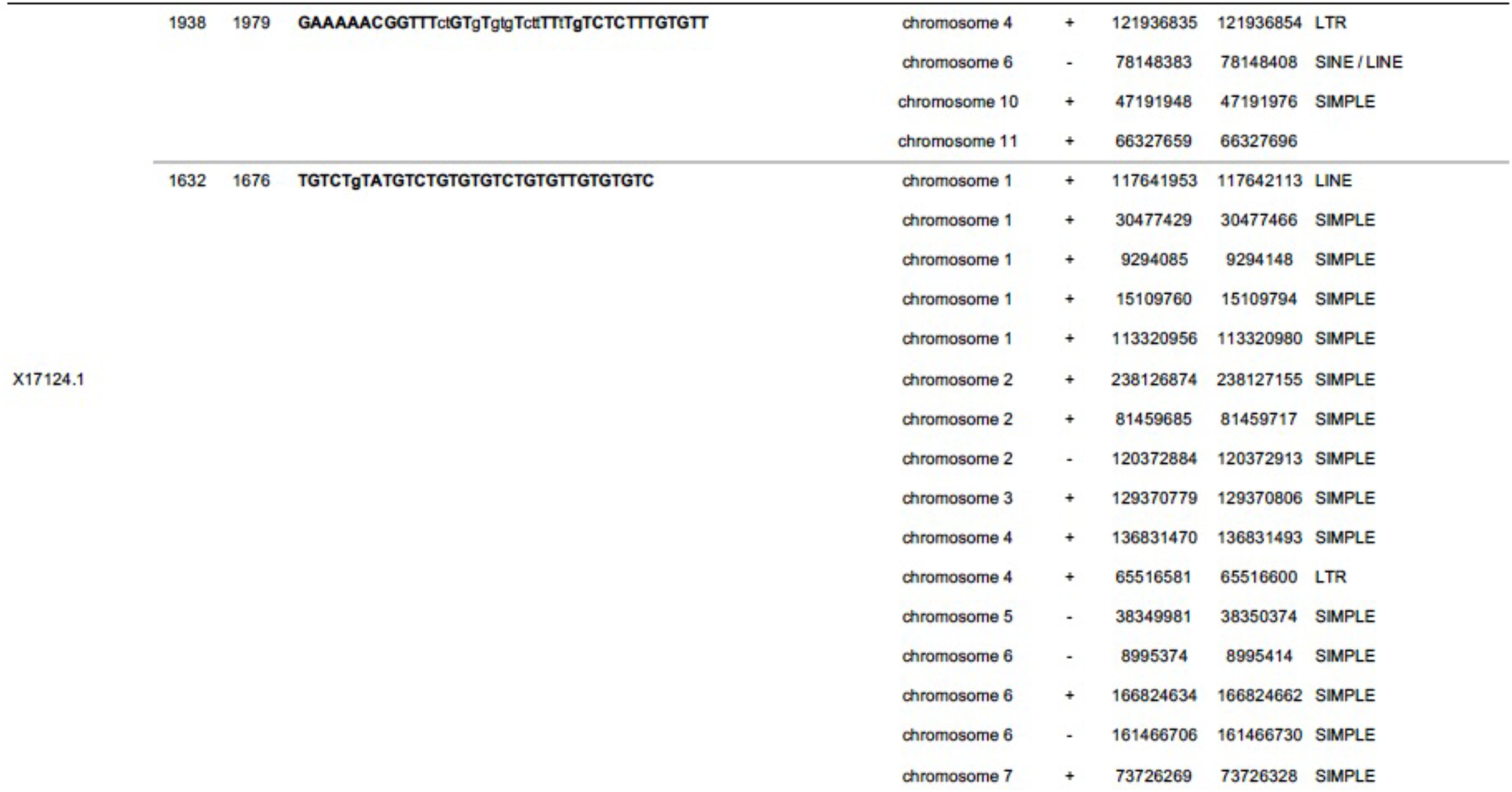
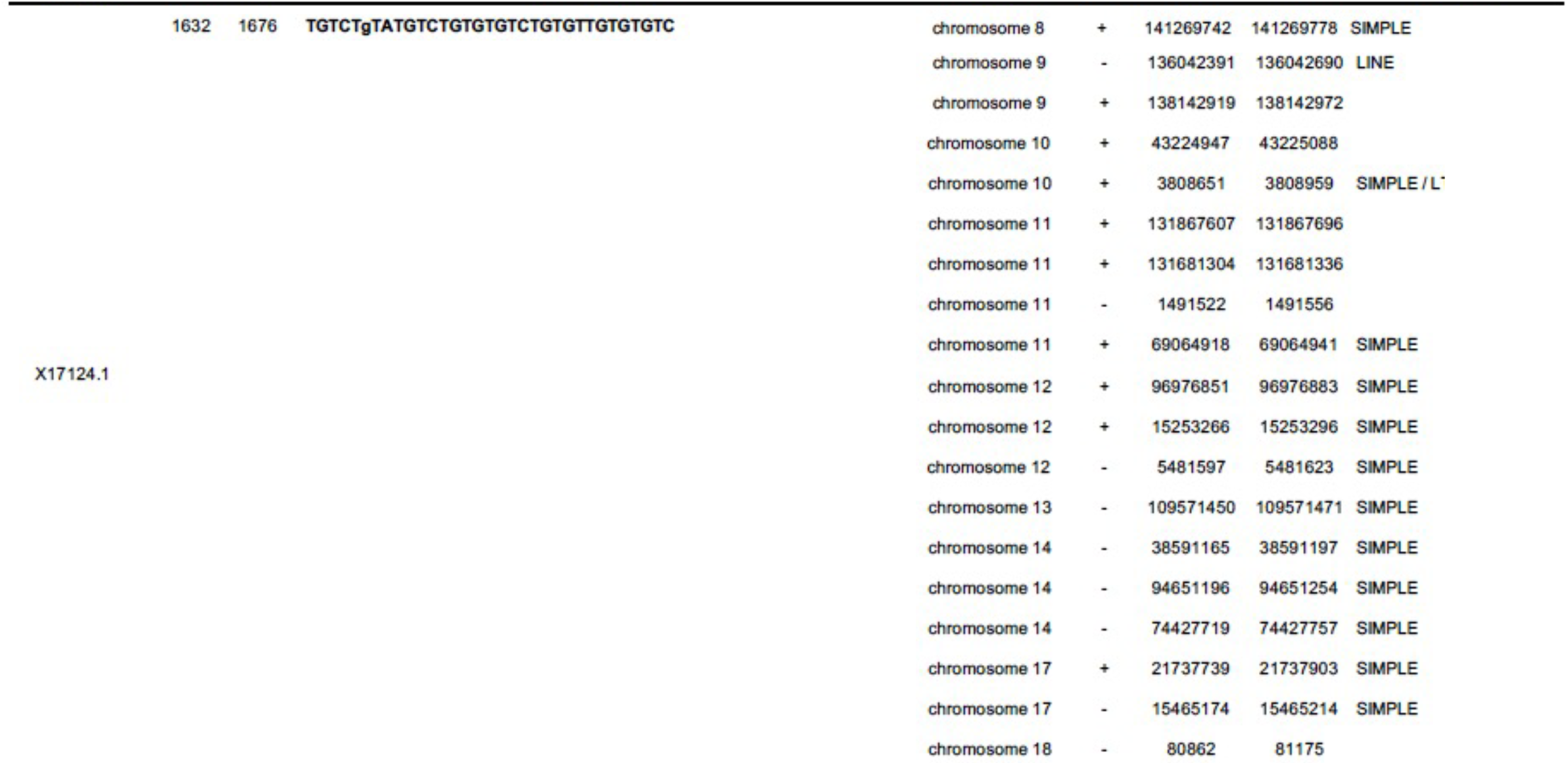
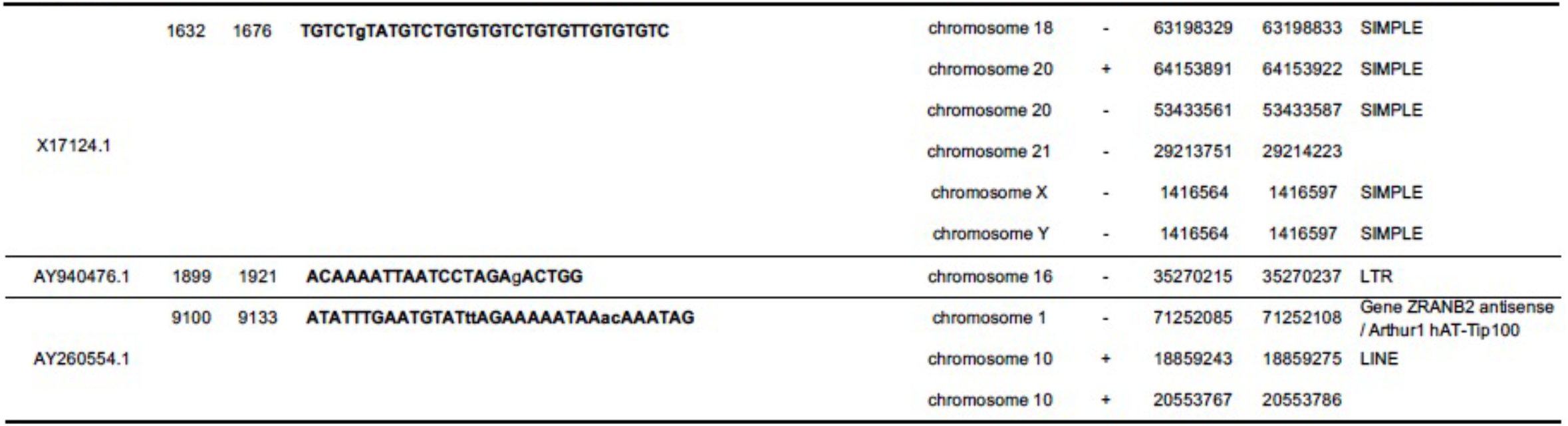
Sequence homologies between various VL30 genomes and the human genome. The Genome Browser of the University of California Santa Cruz Genomic Institute (https://genome.ucsc.edu/) was employed to identify sequence homologies between various VL30 genomes and the human genome. The accession numbers comprising the sequences of the various VL30 genomes, which were employed in the sequence VL30/human homology screen are indicated. The position of the first and last (start and end respectively) VL30 nucleotides in the homologous sequence are shown. VL30 nucleotides that matched their respective nucleotides in homologous human sequences are shown in capital letters. The number of human chromosomes containing sequences with homology to VL30 genome sequences are indicated. The orientations of the relevant homologous strands are shown. The position of the first and last (start and end respectively) nucleotides in the human chromosomes with homology to the relevant VL30 sequence are shown. The repetitive sequences in the human chromosomes with homology to the abovementioned VL30 genomes are defined.

## Discussion

Following successful clinical trials, in 2017 [35, 36] the US Food and Drug Administration (FDA) approved Tisagenlecleucel (Kymriah® Novartis) and axicabtagene ciloleucel (Yescarta® Kite Pharma) as the first two gene therapy-based immunotherapy drugs for patients with hematologic malignancies. The active component of the novel anticancer drugs is made of autologous T-Cells that express viral vector-delivered chimeric antigen receptors (CARs). The CAR-Ts in both drugs target the B cell CD19 protein. Initially, Kymriah® and Yescarta® were indicated for young (under 25 years old) patients with acute lymphocytic leukemia (ALL) or adult patients with large B-cell lymphoma respectively, who relapsed or did not respond to two conventional anticancer treatments. [35, 36] HIV-1-based vectors generated by transient four-plasmid transfection of human cells deliver the CAR expression cassette to Kymriah® CAR-Ts. Generation of Yescarta® CAR-Ts is mediated by transduction with the γ-RVV PG13-CD19-H3. The vector is produced in a stable producer cell line, which was isolated by Kochenderfer *et al*. [5] as a single cell clone (clone H3) following transduction of the murine PG13 packaging cells [6] (ATCC CRL-10686). Prior to and since 2017, the PG13 packaging cell line and its derivatives have been employed to generate γ-RVV-transduced T-cells for various anticancer protocols. [37-46]. Importantly, all tested γ-RVVs employed in the abovementioned clinical trials were determined to be free of replication-competent retroviruses (RCRs). [38, 40] In 2020, the FDA and the European Medicines Agency (EMA) approved a second medicine premised on γ-RVV-transduced CAR-T cells, Tecratus, as an immunotherapy medicine. [47, 48] Tecratus is indicated for the treatment of adult patients with relapsed/refractory mantle cell lymphoma (MCL). The γ-RVVs employed to generate Tecratus and Yescarta® CAR-T cells are identical and are generated in the same PG13 producer cell line. Although stable packaging cell lines facilitate production the of RVVs [7], the transcriptional activity and replication of endogenous murine retrotransposons [8-10, 49] raise biosafety concerns regarding co-packaging of endogenous murine retrotransposons along with clinical-grade γ-RVVs. Indeed, earlier pre-clinical studies reported on efficient co-packaging of the murine VL30 retrotransposon with γ-RVVs generated in various murine stable packaging cells. [11-15, 17] VL30 is a nonautonomous LTR-retrotransposon. Markopoulos *et al*. [50] identified 372 VL30 sequences in the mouse genome. The murine VL30 sequences include 86 full-length genomes comprising 2 LTRs and all the *cis* elements required for retroviral replication, including a primer-binding site (PBS), a packaging signal and a polypurine tract (PPT). VL30 promoter/enhancer sequences in the 5’ LTR contain multiple transcription factor binding sites, which regulate VL30 mRNA expression and potentially exert transcriptional *cis* effects on neighboring host genes. [49-51] VL30 genomes do not encodes functional proteins. [19] Thus, the entire VL30 life cycle is dependent on either exogenous or endogenous retroviral proteins. [19, 50-52] However, VL30 mRNAs function as lncRNAs, most of which contain two RNA motifs, which efficiently bind and alter interaction of the human polypyrimidine tract-binding protein-associated splicing factor (PSF) with its natural DNA and RNA target sites. [20-22] PSF belongs to the Drosophila behavior/human splicing (DBHS) protein family and interacts with nuclear and cytoplasmic proteins, as well as with RNA and DNA target sequences. Nucleic acids/PSF interactions are mediated by a DNA binding domain (DBD) and two RNA binding domains in the PSF protein. [26, 53] PSF is a multifunctional protein involved in major physiological and pathological pathways, including oncogenesis, DNA repair, RNA processing, cytokine release, viral infection, and neurodegeneration [20, 26, 53-56] In mammalian cells, PSF is one of three proteins whose association with the scaffold lncRNA NEAT1_2 initiates a liquid-liquid phase separation process and the formation of membrane-less nuclear paraspeckles organelles, which regulate gene expression via several mechanisms, including sequestration of nucleoplasmic proteins and RNA molecules. An increase in paraspeckle number or size secondary to overexpression of the NEAT1_2 mRNA further recruits paraspeckles proteins, diminishes their nucleoplasmic concentration, and alters their ability to regulate the expression of genes involved in major physiological pathways.[57, 58] VL30 and NEAT1_2 are not the only RNA molecules with which PSF interacts. A study by Song *et al*. demonstrated PSF-binding to four human mRNAs, which are overexpressed in cancer cells. [20] The ability of PSF/VL30 mRNA interaction to promote metastasis of human melanoma cells in immunodeficient mice indicated on the oncogenic potential of VL30 and other PSF-binding ncRNAs. Garen *et al*. outlined a molecular model of ncRNA/tumor suppressor protein (TSP) complex-induced tumorigenesis. [59, 60] In this model, PSF-like TSPs bind to and inhibit transcription of protooncogenes. Binding of PSF to lncRNAs, such as VL30, results in PSF dissociation from its genomic target sequences and consequent activation of transcriptionally suppressed protooncogenes. However, various experimental systems demonstrated that the direction and mechanisms by which PSF/ncRNA complexes affect oncogenic pathways are cell type-dependent. [61-65] Similarly, notwithstanding the role of NEAT1 and PSF in inflammation and activation of the innate immune response [24, 66-69], paraspeckles’ effects on viral infection are pathogen-specific. [24, 57, 64, 70-75] Based on the multiple mechanism by which PSF mediates its functions, it is difficult to predict the effects of VL30 mRNA expression on inflammatory (e.g., cytokine release) or oncogenic pathways in human T-cells. However, the oncogenic potential of PSF-binding mRNAs was considered in the wake of an earlier gene therapy pre-clinical study demonstrating the presence of VL30 genomes in lymphoma cells following bone marrow transplantation of γ-RVV-transduced simian hematopoietic stem cells. [16] In this study, the abovementioned VL30 genome containing lymphoma cells did not express VL30 mRNAs, which suggested that additional mechanisms of VL30-mediated insertional mutagenesis can contribute to the oncogenic potential of VL30 genomes. Reverse-transcribed VL30 genomes preferentially integrate in proximity to transcriptional start sites, and their distribution among mouse chromosomes is not random. [50] VL30-mediated insertional mutagenesis can alter host gene expression via various mechanisms, including a) directly disrupting of host regulatory and protein-encoding sequences, b) transcriptionally activating of neighboring host genes via transcription factor-binding sites in the VL30 LTRs [50, 51, 76-78], and c) spreading host-mediated epigenetic silencing from the VL30 LTR to neighboring genes. [79-81] The integration of transcriptionally active VL30 genomes into patients’ chromatin raises an additional biosafety concern regarding the possibility of emerging novel retroviruses following recombination between VL30 genomes and either endogenous or exogenous retroviruses. Genomic analysis of the transforming and replication-defective Kirsten and Harvey murine sarcoma viruses (Ki-MSV and Ha-MSV, respectively) hints at the recombinational potential of VL30. Evolved by a series of recombination events. The genomes of these viruses comprise sequences from three sources: the rat VL30, the Moloney murine leukemia virus LTRs, and the Kirsten and Harvey viral *ras* genes, respectively. [33, 34, 82] In a different study, Itin *et al*. identified DNA recombinants comprising the VL30 LTRs and sequences with homology to the Murine Leukemia Virus (MuLV) *gag* and *pol* genes. [28] Several VL30-specific factors potentially contribute to the risk of emerging novel VL30-based recombinants, these factors include a) the presence of multiple short sequences in the human genome that exhibit high identity to sequences in VL30 genomes and b) packaging of VL30 genomes into productive human retroviral particles. Under natural conditions, sequence and structural differences between *cis* regulatory elements involved in all the steps of the human retroviral (e.g., HIV-1and HTLV) and VL30 life cycle minimize the risk of horizontal transfer of murine VL30 genomes by HIV-1 particles. For instance, there is no homology between the PBS sequences of VL30 strains (which are mostly complementary to the t-RNA^Gly^) and the sequence of the HIV-1 PBS (which is complementary to the t-RNA^Lys^). Similarly, the 3’ polypurine tract (PPT) of the VL30 shows no sequence homology with that of the HIV-1. [18, 19, 49, 50, 83] Furthermore, there is no sequence homology between the VL30 and the HIV-1 packaging signals. Notwithstanding the lack of sequence homology between key VL30 and the HIV-1 *cis* elements, in this study, for the first time, we showed reverse transcription-dependent delivery of transcriptionally active VL30 genomes to naïve human cells by HIV-1 vector particles. However, the loss of HIV-1 vector-delivered VL30 genomes in replicating cells (following 4 passages in culture) suggested that reverse-transcribed VL30 DNA failed to integrate into transduced cells’ chromatin. This phenomenon can be attributed to incompatibility between the VL30 *att* sites and the HIV-1 integrase [29] or/and to incompatibility between the VL30 polypurine tract (PPT) and the HIV-1 reverse transcriptase, which may alter the last step of reverse transcription and consequently leads to the formation of mostly episomal single-LTR circles (which, unlike fully reverse-transcribed linear double-stranded genomes, cannot serve as an integration template).[84] This notion is supported by an earlier study demonstrating the delivery of productive γ-retroviral vectors by HIV-1 particles which, similar to the abovementioned VL30 genomes, failed to integrate. In this study, genomic analysis of γ-retroviral vector genomes (following delivery by HIV-1 vector particles) demonstrated the presence of episomal single-LTR circles and no linear vector forms. [30] Importantly, incorporation of HIV sequences comprising rev-response element (RRE) to γ-retroviral vector genomes significantly increased their titers following packaging by HIV-1 particles. Premised on these findings, as well as the natural reversion rate of mutated HIV-1 genomes [85, 86] and the VL30 recombinational potential [28, 33, 34], it is possible to speculate that in the presence of active HIV-1 genomes, novel VL30 recombinants can evolve to maximize their transfer by HIV-1 particles. This theoretical scenario raises biosafety concerns regarding mobilization of VL30 genomes within CAR-T treated patients and in the worst scenario within the treated patients’ community. Additional consideration should be given to the fact that PFS-mediated alternative splicing is central to T-cell activation. Thus, there is a possibility that CAR-T cell activation (via the CAR CD28-costimulatory domain) may be altered following binding/sequestration of nuclear PFS by VL30 mRNA. [87-89]

Notwithstanding these biosafety concerns, more than a thousand patients with incurable oncologic diseases have been successfully treated with CAR-T cells. [90, 91] To date, not a single clinical report has described proliferative abnormalities that could be attributed to the inadvertent contamination of therapeutic T-cells with VL30 genomes. [38, 40, 91] Theoretically, it is remotely possible that in contrast to the PG13 producer cells characterized in this study, specific vector producer cell clones that were individually isolated for specific clinical applications did not secrete VL30 genomes-comprising γ-retroviral particles. Furthermore, this study did not characterize the presence of VL30 genomes in patient-administered CAR-T cells. Thus, the level of VL30 mRNA expression in patient-transplanted T-cells and the half-life of VL30 genome-containing T-cells has not been evaluated. These unknown variables and the clinical history of a large number of CAR-T cells-treated patients suggest that there are no imminent biosafety risks associated with potentially VL30 genome-containing CAR-T cells. Importantly, short- and long-term biosafety concerns associated with the production of therapeutic γ-retroviral vector carrying CAR-T cells in murine packaging cell lines could be avoided by using human packaging cell lines to generate the same CAR-carrying γ-retroviral vectors. Importantly, this study underscores the importance of addressing the potential biosafety risks associated with the transfer of non-human endogenous retroviruses and potentially nonautonomous retrotransposons to patients undergoing novel therapeutic procedures.[92-96]

## Materials and Methods

### Cells

The PG13 γ-retroviral vector packaging cell line was purchased from the American Type Culture Collection (CRL-10686, ATCC, Manassas, VA). PG13 and 293T cells were cultured in Dulbecco’s Modified Eagle’s Medium (DMEM)-high glucose (10-013-CV, Corning, Mediatech, Inc., Manassas, VA) with 1x antibiotic-antimycotic (15240-062, Gibco, Thermo Fisher Scientific, Grand Island, NY) and 10% fetal bovine serum (10437-028, Gibco, Thermo Fisher Scientific, Grand Island, NY).

Peripheral blood mononuclear cells (PBMCs) were isolated from buffy coats (Gulf Coast Regional Blood Center) using Lymphoprep (AN1001967, Accurate Chemical and Scientific Corporation, Carle Place, NY). Next PBMCs were then activated using 1 µg/mL immobilized CD3 and CD28 antibodies (Miltenyi Biotec, Cambridge, MA). After 3 days, cells were transduced in 24-well plates precoated with recombinant fibronectin (FN CH-296, Retronectin Takara, Clontech, Mountain View, CA) with retroviral supernatants and expanded in complete medium (45% RPMI-1640 and 45% Click’s medium, 10% FBS, and 2 mM GlutaMAX) supplemented with IL-7 (10 ng/mL, Miltenyi) and IL-15 (5 ng/mL, Miltenyi). After transduction, T-cells were expanded *ex vivo* in complete medium (45% RPMI-1640 and 45% Click’s medium, 10% FBS, and 2 mM GlutaMax) in the presence of IL-7 (10 ng/ml) and IL-15 (5 ng/ml), which were added twice a week for up to 15 days.

### Isolation of single-cell 293T clones containing VL30 genomes

293T cells were transduced with the lentiviral vector pTK1261, from which the firefly luciferase cDNA and the fusion GFP/blasticidin marker gene were expressed under the control of a CMV promoter and the encephalomyelitis virus internal ribosome entry site (IRES), respectively. Transduced cells were selected for blasticidin resistance in the presence of 50µg/ml blasticidin (293T-1261). Blasticidin resistant 293T cells were co-cultured with PG13 cells. At confluency, the co-culture (293T-1261/PG13) cell population was passaged, and PG13 cells were eliminated in the presence of 50µg/ml blasticidin. The abovementioned process of co-culturing was repeated seven times, after which single cell clones of blasticidin-resistant 293T cells were isolated. The absences of contaminating PG13 cells was confirmed by the lack of PCR amplification of the mouse GAPDH gene. GCN in the abovementioned isolated single-cell clones was determined by qPCR as described below.

### Production CAR carrying γ-RVV from a newly established stable producer cell line

A stock of Eco-pseudotyped SFG.iC9.GD2.CAR.IL15 γ–RVV was produced by transient transfection of the ecotropic ΦNX-Eco packaging cell line (American Type Culture Collection product CRL-3214; ATCC, Manassas, VA) with the vector-expression cassette. A heterogenous stable producer cell line was generated by repeated transduction of the murine gibbon ape leukemia virus (GalV) envelope-expressing PG13 packaging cell line (# CRL-10686™, ATCC, Manassas, VA) with the abovementioned Eco-pseudotyped SFG.iC9.GD2 vector. To enrich for stable producer cells with high VCN, the abovementioned heterogenous population of vector producing cells was immuno-stained with an anti-idiotype antibody specific for the GD2 CAR [31] high transgene expressing cells were sorted using the BD Jazz cell sorter. Following limiting dilution single-cell clones of vector-producing cells were isolated and functionally screened for the highest biological titer by qPCR. The highest vector producing clone was expanded for generation of the Master Cell Bank (MCB) for production of GalV envelope-pseudotyped retroviral particles carrying the iC9.GD2.CAR.IL15 expression cassette. For each lot of γ–RVV supernatant, cells from the abovementioned MCB were thawed and expanded in the iCellis Nano bench-top bioreactor, (Pall, Inc). The iCellis Nano is a fixed bed bioreactor using fiber material for the attachment of the attachment of adherent vector producer cells. The system was employed to provide continuous monitoring of dissolved oxygen, pH, and temperature. The system provided continuous circulation of culture media along with proper gas mixture of O_2_ and CO_2_. pH was additionally controlled by the addition of base as required. After seeding onto the fiber bed of the iCellis Nano, the cells are cultured for 2 - 3 days to reach near confluency. At this point, the media was replaced with fresh media, and following overnight culture, the supernatant was harvested into a transfer bag, and fresh media was added to the system. This supernatant is termed “Day 1” harvest. This process was repeated four additional times resulting in a total of five days of supernatant harvest. After harvest of the fifth supernatant, the producer cells were removed from the bioreactor using enzymatic digestion (TrypLE) and washing with PBS. The harvested cells were counted and samples taken for QC studies. The MCB, end of production cells and viral supernatant were tested following FDA recommendation for sterility and absence of replication competent retroviral particles.

### Lentiviral and γ-retroviral vectors

The construction of the non-SIN γ-RVV, SFG.iC9.GD2.CAR.IL15 was described earlier. [31] pTK1261 was constructed by cloning a BglII/BamHI DNA fragment containing the firefly luciferase cDNA into a BamHI site in the lentiviral vector pTK642.[97] pTK2229 was constructed by cloning an AfeI/HpaI DNA fragment comprising a VL30 sequence from nucleotides 2140 to 2769 of GeneBank: AF486451.1 into a PshAI site in a lentiviral vector comprising a CMV promoter-regulated expression cassette (encoding the mCherry-T2A-Puromycin selection marker) in opposite orientation to the LTRs.

The pTK1808 and pTK2151 vectors were purchased from Addgene (Addgene, Watertown, Cat# 14088 and 10668, respectively). Both vectors are premised on the γ-retroviral vector pBabe-puro and carry either the SV40 large-T antigen or the green fluorescence protein (GFP) cDNA under the control of the vector 5’ LTR, respectively. Note that downstream to the SV40 large-T antigen cDNA, the pTK1808 vector also contains the puromycin resistance cDNA under the control of the SV40 promoter.

### Production of viral vectors by transient transfection

All vector particles were VSV-g pseudotyped and produced in 293T cells using the transient three-plasmid calcium phosphate transfection method as described earlier. [98, 99] In brief, the second-generation lentiviral vector packaging cassette ΔNRF [100] or the MLV Gag/Pol expression cassette (a kind gift from Dr. Nikunj Somia at the University of Minnesota) were transiently transfected into 293T cells along with the VSV-G envelope expression cassette and the relevant vector construct. Vector particles in conditioned media were harvested ∼60 hours after transfection and filtered through a 0.45-µm syringe filter.

### Analysis of VL30 genome- and γ-retroviral vector copy number (GCN and VCN, respectively)

Genomic DNA samples were extracted by the DNeasy Blood & Tissue Kit (69506, QIAGEN, Hilden, Germany). To eliminate potential contamination with carried-over transfected plasmid DNA, all genomic DNA samples (except DNA samples extracted from cultured human T-cells) were digested with the DpnI restriction enzyme.

GCN and VCN were determined by quantitative real-time polymerase chain reaction (qPCR) using the QuantStudio™ 3 system (Applied Biosystems, Thermo Fisher Scientific, Waltham, MA). To measure VL30 GCN, a reference cell population containing a single-copy of a VL30 sequence was established (293T-2229 cells). Specifically, 293T cells were transduced with the lentiviral vector pTK2229 (at MOI <0.001) and selected for puromycin resistance in the presence of 5 µg/ml of puromycin (P8833, MilliporeSigma, St. Louis, MO). DNA samples extracted from the abovementioned heterogenous population of puromycin resistant 293T cells served to establish a reference DNA standard-curve to measure VL30 VCN by qPCR using a primer/probe set (Integrated DNA Technologies, Inc., Coralville, Iowa) comprising a forward primer 5’-CCTTGACCAGAAGCCACTATG-3’, a reverse primer 5’-TCAGAGATTGGGACCCTGAA-3’, and a 6-FAM™-conjugated probe 5’-TGTAAGATGGCCTGCTTGT CTGCA-3’.

To measure the VCN of γ-RVVs (expressing either a CAR or the GFP cDNA), a reference cell population comprising a single-copy of a γ-RVV genome was established (293T-1808 cells). Specifically, 293T cells were transduced with the γ-RVV l vector pTK1808 (at MOI <0.001) and selected for puromycin resistance in the presence of 5 µg/ml of puromycin (MilliporeSigma, St. Louis, MO). DNA samples extracted from the abovementioned heterogenous population of puromycin-resistant 293T cells served to establish a reference DNA standard-curve to measure γ–RVV VCN by qPCR using a primer/probe set (Integrated DNA Technologies, Inc., Coralville, Iowa) comprising a forward primer 5’-CGCTGACGGGTAGTCAATC-3’, a reverse primer 5’-GGGTACCCGTGTATCCAATAAA-3’ and a 6-FAM™ probe 5’-ACTTGTGGTCTCGCTGTTCCTTGG-3’. qPCR analysis of the endogenous human RNaseP, which served as an internal reference control, was premised on a commercial human RNaseP primer/probe set (443328, Applied Biosystems, Thermo Fisher Scientific, Waltham, MA).

PCR amplification of the mouse GAPDH gene was premised on a primer/probe set from Hoffmann-La Roche Ltd, Basel, Switzerland (Universal ProbeLibrary Mouse GAPD Gene Assay, 05046211001). qPCR was performed with the ABsolute qPCR ROX Mix (AB-1138/B, Applied Biosystems, Thermo Fisher Scientific, Waltham, MA) under the following conditions: 95°C for 15 min, and then 40 cycles of 95°C for 15 sec and 60°C for 1 min.

### RNA isolation and qRT-PCR

RNA was isolated using an RNeasy® Plus Mini Kit (QIAGEN, Hilden, Germany) and converted to cDNA using a QuantiTect® Reverse Transcription Kit (QIAGEN, Hilden, Germany). qRT-PCR of VL30 and γ-RVV mRNA was premised on the same primer/probe sets that were described above. The qRT-PCR assay for the human ACTB mRNA, which served as an internal reference control, was premised on a commercial set of primers/probe (Hs.PT.39a.22214847) conjugated with HEX™ at the 5’ end (Integrated DNA Technologies, Inc., Coralville, Iowa). All PCR results were analyzed with Prism 9 software (GraphPad Software, San Diego, CA).

### Statistical analysis

VL30 genome copy number (GCN) and γ-RVV vector copy number (VCN) in human T-cells transduced with CAR-expressing γ-RVV were calculated as the average of 3 technical replicates. The association between VL30 GCN and γ-RVV in primary human CAR-T cells was estimated by Pearson correlation and tested with linear regression analysis. Statistical analyses employed to characterize the significance of various treatments’ effects on VL30 GCN and γ-RVV VCN are outlined in the figure legends.

## Disclosure

TK is an inventor of PPT-deleted lentiviral vectors and of integration defective lentiviral vector production technologies, which are owned by the University of North Carolina. Some of these technologies are licensed to a commercial entity. GD is a paid consultant for Bellicum Pharmaceuticals, Tessa Therapeutics and Catamaran. BS is supported by Bluebirdbio, Bellicum Pharmaceutical, Cell Medica, Tessa therapeutics and is a paid consultant for Tessa Therapeutics. The other authors declare that they have no conflict of interest.

## Acknowledgements

The following reagents were obtained through the National Institutes of Health (NIH) AIDS Research and Reference Reagent Program, Division of AIDS, the National Institute of Allergy and Infectious Diseases: the HIV-1 p24 monoclonal antibody (183-H12-5C) from Bruce Chesebro and Kathy Wehrly. The study was supported by NIH grants R01-HL128119 (to SHL and TK), R01-DK058702 (to SHL and TK). This study is dedicated to the US Marine Corps and the Gold Star families. In memory of Henryk Goldszmit and Stanislav Petrov.

## Notes

### Summary of Updates

Adding conflicts of interest

## References

1. FDA Approves Second CAR T-cell Therapy. Cancer Discov, 2018. 8(1): p. 5–6.

2. Locke, F.L., W.Y. Go, and S.S. Neelapu, Development and Use of the Anti-CD19 Chimeric Antigen Receptor T-Cell Therapy Axicabtagene Ciloleucel in Large B-Cell Lymphoma: A Review. JAMA Oncol, 2020. 6(2): p. 281–290.

3. Locke, F.L., et al., Phase 1 Results of ZUMA-1: A Multicenter Study of KTE-C19 Anti-CD19 CAR T Cell Therapy in Refractory Aggressive Lymphoma. Mol Ther, 2017. 25(1): p. 285–295.

4. Hughes, M.S., et al., Transfer of a TCR gene derived from a patient with a marked antitumor response conveys highly active T-cell effector functions. Hum Gene Ther, 2005. 16(4): p. 457–72.

5. Kochenderfer, J.N., et al., Construction and preclinical evaluation of an anti-CD19 chimeric antigen receptor. J Immunother, 2009. 32(7): p. 689–702.

6. Miller, A.D., et al., Construction and properties of retrovirus packaging cells based on gibbon ape leukemia virus. J Virol, 1991. 65(5): p. 2220–4.

7. Miller, A.D., Retrovirus packaging cells. Hum Gene Ther, 1990. 1(1): p. 5–14.

8. Gagnier, L., V.P. Belancio, and D.L. Mager, Mouse germ line mutations due to retrotransposon insertions. Mob DNA, 2019. 10: p. 15.

9. Huang, C.R., K.H. Burns, and J.D. Boeke, Active transposition in genomes. Annu Rev Genet, 2012. 46: p. 651–75.

10. Maksakova, I.A., et al., Retroviral elements and their hosts: insertional mutagenesis in the mouse germ line. PLoS Genet, 2006. 2(1): p. e2.

11. Scadden, D.T., B. Fuller, and J.M. Cunningham, Human cells infected with retrovirus vectors acquire an endogenous murine provirus. J Virol, 1990. 64(1): p. 424–7.

12. Chakraborty, A.K., M.A. Zink, and C.P. Hodgson, Transmission of endogenous VL30 retrotransposons by helper cells used in gene therapy. Cancer Gene Ther, 1994. 1(2): p. 113–8.

13. Hatzoglou, M., et al., Efficient packaging of a specific VL30 retroelement by psi 2 cells which produce MoMLV recombinant retroviruses. Hum Gene Ther, 1990. 1(4): p. 385–97.

14. Howk, R.S., et al., Identification of a 30S RNA with properties of a defective type C virus in murine cells. J Virol, 1978. 25(1): p. 115–23.

15. Patience, C., et al., Packaging of endogenous retroviral sequences in retroviral vectors produced by murine and human packaging cells. J Virol, 1998. 72(4): p. 2671–6.

16. Purcell, D.F., et al., An array of murine leukemia virus-related elements is transmitted and expressed in a primate recipient of retroviral gene transfer. J Virol, 1996. 70(2): p. 887–97.

17. Sherwin, S.A., et al., Rescue of endogenous 30S retroviral sequences from mouse cells by baboon type C virus. J Virol, 1978. 26(2): p. 257–64.

18. Song, X., et al., Retroviral-mediated transmission of a mouse VL30 RNA to human melanoma cells promotes metastasis in an immunodeficient mouse model. Proc Natl Acad Sci U S A, 2002. 99(9): p. 6269–73.

19. Adams, S.E., et al., Complete nucleotide sequence of a mouse VL30 retro-element. Mol Cell Biol, 1988. 8(8): p. 2989–98.

20. Li, L., et al., Role of human noncoding RNAs in the control of tumorigenesis. Proc Natl Acad Sci U S A, 2009. 106(31): p. 12956–61.

21. Song, X., A. Sui, and A. Garen, Binding of mouse VL30 retrotransposon RNA to PSF protein induces genes repressed by PSF: effects on steroidogenesis and oncogenesis. Proc Natl Acad Sci U S A, 2004. 101(2): p. 621–6.

22. Wang, G., et al., Regulation of proto-oncogene transcription, cell proliferation, and tumorigenesis in mice by PSF protein and a VL30 noncoding RNA. Proc Natl Acad Sci U S A, 2009. 106(39): p. 16794–8.

23. Hadjicharalambous, M.R. and M.A. Lindsay, Long Non-Coding RNAs and the Innate Immune Response. Noncoding RNA, 2019. 5(2).

24. Imamura, K., et al., Long noncoding RNA NEAT1-dependent SFPQ relocation from promoter region to paraspeckle mediates IL8 expression upon immune stimuli. Mol Cell, 2014. 53(3): p. 393–406.

25. Morchikh, M., et al., HEXIM1 and NEAT1 Long Non-coding RNA Form a Multi-subunit Complex that Regulates DNA-Mediated Innate Immune Response. Mol Cell, 2017. 67(3): p. 387–399 e5.

26. Yarosh, C.A., et al., PSF: nuclear busy-body or nuclear facilitator? Wiley Interdiscip Rev RNA, 2015. 6(4): p. 351–67.

27. Zhou, B., et al., Endogenous Retrovirus-Derived Long Noncoding RNA Enhances Innate Immune Responses via Derepressing RELA Expression. mBio, 2019. 10(4).

28. Itin, A. and E. Keshet, Apparent recombinants between virus-liKE (VL30) and murine leukemia virus-related sequences in mouse DNA. J Virol, 1983. 47(1): p. 178–84.

29. Nightingale, S.J., et al., Transient gene expression by nonintegrating lentiviral vectors. Mol Ther, 2006. 13(6): p. 1121–32.

30. Cockrell, A.S., et al., The HIV-1 Rev/RRE system is required for HIV-1 5’ UTR cis elements to augment encapsidation of heterologous RNA into HIV-1 viral particles. Retrovirology, 2011. 8: p. 51.

31. Chen, Y., et al., Eradication of Neuroblastoma by T Cells Redirected with an Optimized GD2-Specific Chimeric Antigen Receptor and Interleukin-15. Clin Cancer Res, 2019. 25(9): p. 2915–2924.

32. Coffin, J.M., Structure, replication, and recombination of retrovirus genomes: some unifying hypotheses. J Gen Virol, 1979. 42(1): p. 1–26.

33. Ellis, R.W., et al., The p21 src genes of Harvey and Kirsten sarcoma viruses originate from divergent members of a family of normal vertebrate genes. Nature, 1981. 292(5823): p. 506–11.

34. Roop, D.R., et al., An activated Harvey ras oncogene produces benign tumours on mouse epidermal tissue. Nature, 1986. 323(6091): p. 822–4.

35. Maude, S.L., et al., Chimeric antigen receptor T cells for sustained remissions in leukemia. N Engl J Med, 2014. 371(16): p. 1507–17.

36. Neelapu, S.S., et al., Axicabtagene Ciloleucel CAR T-Cell Therapy in Refractory Large B-Cell Lymphoma. N Engl J Med, 2017. 377(26): p. 2531–2544.

37. Albinger, N., J. Hartmann, and E. Ullrich, Current status and perspective of CAR-T and CAR-NK cell therapy trials in Germany. Gene Ther, 2021. 28(9): p. 513–527.

38. Bear, A.S., et al., Replication-competent retroviruses in gene-modified T cells used in clinical trials: is it time to revise the testing requirements? Mol Ther, 2012. 20(2): p. 246–9.

39. Brentjens, R.J., et al., Safety and persistence of adoptively transferred autologous CD19-targeted T cells in patients with relapsed or chemotherapy refractory B-cell leukemias. Blood, 2011. 118(18): p. 4817–28.

40. Cornetta, K., et al., Screening Clinical Cell Products for Replication Competent Retrovirus: The National Gene Vector Biorepository Experience. Mol Ther Methods Clin Dev, 2018. 10: p. 371–378.

41. Davila, M.L., et al., Efficacy and toxicity management of 19-28z CAR T cell therapy in B cell acute lymphoblastic leukemia. Sci Transl Med, 2014. 6(224): p. 224ra25.

42. Hollyman, D., et al., Manufacturing validation of biologically functional T cells targeted to CD19 antigen for autologous adoptive cell therapy. J Immunother, 2009. 32(2): p. 169–80.

43. Itzhaki, O., et al., Head-to-head comparison of in-house produced CD19 CAR-T cell in ALL and NHL patients. J Immunother Cancer, 2020. 8(1).

44. Pule, M.A., et al., Virus-specific T cells engineered to coexpress tumor-specific receptors: persistence and antitumor activity in individuals with neuroblastoma. Nat Med, 2008. 14(11): p. 1264–70.

45. Ramos, C.A., et al., Clinical and immunological responses after CD30-specific chimeric antigen receptor-redirected lymphocytes. J Clin Invest, 2017. 127(9): p. 3462–3471.

46. Ramos, C.A., et al., Clinical responses with T lymphocytes targeting malignancy-associated kappa light chains. J Clin Invest, 2016. 126(7): p. 2588–96.

47. EMA, Meeting highlights from the Committee for Medicinal Products for Human Use (CHMP) 12–15 October 2020.

48. FDA, U.S., Biologics License Application (BLA) for brexucabtagene autoleucel. 2020. p. July 24, 2020 Approval Letter—TECARTUS; Food and Drug Administration: Silver Spring, MD, USA, 2020.

49. French, N.S. and J.D. Norton, Structure and functional properties of mouse VL30 retrotransposons. Biochim Biophys Acta, 1997. 1352(1): p. 33–47.

50. Markopoulos, G., et al., Genomic analysis of mouse VL30 retrotransposons. Mob DNA, 2016. 7: p. 10.

51. Schiff, R., A. Itin, and E. Keshet, Transcriptional activation of mouse retrotransposons in vivo: specific expression in steroidogenic cells in response to trophic hormones. Genes Dev, 1991. 5(4): p. 521–32.

52. Besmer, P., et al., Virus-like 30S RNA in mouse cells. J Virol, 1979. 29(3): p. 1168–76.

53. Lim, Y.W., et al., The Emerging Role of the RNA-Binding Protein SFPQ in Neuronal Function and Neurodegeneration. Int J Mol Sci, 2020. 21(19).

54. Gordon, P.M., et al., A conserved role for the ALS-linked splicing factor SFPQ in repression of pathogenic cryptic last exons. Nat Commun, 2021. 12(1): p. 1918.

55. Ji, Q., et al., Long non-coding RNA MALAT1 promotes tumour growth and metastasis in colorectal cancer through binding to SFPQ and releasing oncogene PTBP2 from SFPQ/PTBP2 complex. Br J Cancer, 2014. 111(4): p. 736–48.

56. Takayama, K.I., et al., Dysregulation of spliceosome gene expression in advanced prostate cancer by RNA-binding protein PSF. Proc Natl Acad Sci U S A, 2017. 114(39): p. 10461–10466.

57. Fox, A.H., et al., Paraspeckles: Where Long Noncoding RNA Meets Phase Separation. Trends Biochem Sci, 2018. 43(2): p. 124–135.

58. Hirose, T., et al., NEAT1 long noncoding RNA regulates transcription via protein sequestration within subnuclear bodies. Mol Biol Cell, 2014. 25(1): p. 169–83.

59. Garen, A., From a retrovirus infection of mice to a long noncoding RNA that induces proto-oncogene transcription and oncogenesis via an epigenetic transcription switch. Signal Transduct Target Ther, 2016. 1: p. 16007.

60. Garen, A. and X. Song, Regulatory roles of tumor-suppressor proteins and noncoding RNA in cancer and normal cell functions. Int J Cancer, 2008. 122(8): p. 1687–9.

61. Bi, O., et al., SFPQ promotes an oncogenic transcriptomic state in melanoma. Oncogene, 2021. 40(33): p. 5192–5203.

62. Dong, P., et al., Long Non-coding RNA NEAT1: A Novel Target for Diagnosis and Therapy in Human Tumors. Front Genet, 2018. 9: p. 471.

63. Mello, S.S. and L.D. Attardi, Neat-en-ing up our understanding of p53 pathways in tumor suppression. Cell Cycle, 2018. 17(13): p. 1527–1535.

64. Pisani, G. and B. Baron, Nuclear paraspeckles function in mediating gene regulatory and apoptotic pathways. Noncoding RNA Res, 2019. 4(4): p. 128–134.

65. Zeng, C., et al., The c-Myc-regulated lncRNA NEAT1 and paraspeckles modulate imatinib-induced apoptosis in CML cells. Mol Cancer, 2018. 17(1): p. 130.

66. Menon, M.P. and K.F. Hua, The Long Non-coding RNAs: Paramount Regulators of the NLRP3 Inflammasome. Front Immunol, 2020. 11: p. 569524.

67. Zhang, F., et al., Identification of the long noncoding RNA NEAT1 as a novel inflammatory regulator acting through MAPK pathway in human lupus. J Autoimmun, 2016. 75: p. 96–104.

68. Zhang, M., et al., Knockdown of NEAT1 induces tolerogenic phenotype in dendritic cells by inhibiting activation of NLRP3 inflammasome. Theranostics, 2019. 9(12): p. 3425–3442.

69. Zhang, P., et al., The lncRNA Neat1 promotes activation of inflammasomes in macrophages. Nat Commun, 2019. 10(1): p. 1495.

70. Bu, F.T., et al., LncRNA NEAT1: Shedding light on mechanisms and opportunities in liver diseases. Liver Int, 2020. 40(11): p. 2612–2626.

71. Lee, N., et al., EBV noncoding RNA EBER2 interacts with host RNA-binding proteins to regulate viral gene expression. Proc Natl Acad Sci U S A, 2016. 113(12): p. 3221–6.

72. Liu, H., et al., HIV-1 replication in CD4(+) T cells exploits the down-regulation of antiviral NEAT1 long non-coding RNAs following T cell activation. Virology, 2018. 522: p. 193–198.

73. Wang, Z., et al., NEAT1 modulates herpes simplex virus-1 replication by regulating viral gene transcription. Cell Mol Life Sci, 2017. 74(6): p. 1117–1131.

74. Zhang, Q., et al., NEAT1 long noncoding RNA and paraspeckle bodies modulate HIV-1 posttranscriptional expression. mBio, 2013. 4(1): p. e00596–12.

75. Zolotukhin, A.S., et al., PSF acts through the human immunodeficiency virus type 1 mRNA instability elements to regulate virus expression. Mol Cell Biol, 2003. 23(18): p. 6618–30.

76. Anderson, G.R. and D.L. Stoler, Anoxia, wound healing, VL30 elements, and the molecular basis of malignant conversion. Bioessays, 1993. 15(4): p. 265–72.

77. Eaton, L. and J.D. Norton, Independent regulation of mouse VL30 retrotransposon expression in response to serum and oncogenic cell transformation. Nucleic Acids Res, 1990. 18(8): p. 2069–77.

78. Singh, K., S. Saragosti, and M. Botchan, Isolation of cellular genes differentially expressed in mouse NIH 3T3 cells and a simian virus 40-transformed derivative: growth-specific expression of VL30 genes. Mol Cell Biol, 1985. 5(10): p. 2590–8.

79. Choi, J.Y. and Y.C.G. Lee, Double-edged sword: The evolutionary consequences of the epigenetic silencing of transposable elements. PLoS Genet, 2020. 16(7): p. e1008872.

80. Misiak, B., L. Ricceri, and M.M. Sasiadek, Transposable Elements and Their Epigenetic Regulation in Mental Disorders: Current Evidence in the Field. Front Genet, 2019. 10: p. 580.

81. Sundaram, V. and J. Wysocka, Transposable elements as a potent source of diverse cis-regulatory sequences in mammalian genomes. Philos Trans R Soc Lond B Biol Sci, 2020. 375(1795): p. 20190347.

82. Cox, A.D. and C.J. Der, Ras history: The saga continues. Small GTPases, 2010. 1(1): p. 2–27.

83. Berkhout, B., The primer binding site on the RNA genome of human and simian immunodeficiency viruses is flanked by an upstream hairpin structure. Nucleic Acids Res, 1997. 25(20): p. 4013–7.

84. Kantor, B., et al., Notable reduction in illegitimate integration mediated by a PPT-deleted, nonintegrating lentiviral vector. Mol Ther, 2011. 19(3): p. 547–56.

85. Li, X., et al., Effects of alterations of primer-binding site sequences on human immunodeficiency virus type 1 replication. J Virol, 1994. 68(10): p. 6198–206.

86. Wakefield, J.K., A.G. Wolf, and C.D. Morrow, Human immunodeficiency virus type 1 can use different tRNAs as primers for reverse transcription but selectively maintains a primer binding site complementary to tRNA(3Lys). J Virol, 1995. 69(10): p. 6021–9.

87. Blake, D. and K.W. Lynch, The three as: Alternative splicing, alternative polyadenylation and their impact on apoptosis in immune function. Immunol Rev, 2021. 304(1): p. 30–50.

88. Schaub, A. and E. Glasmacher, Splicing in immune cells-mechanistic insights and emerging topics. Int Immunol, 2017. 29(4): p. 173–181.

89. Topp, J.D., et al., A cell-based screen for splicing regulators identifies hnRNP LL as a distinct signal-induced repressor of CD45 variable exon 4. RNA, 2008. 14(10): p. 2038–49.

90. Crees, Z.D. and A. Ghobadi, Cellular Therapy Updates in B-Cell Lymphoma: The State of the CAR-T. Cancers (Basel), 2021. 13(20).

91. June, C.H. and M. Sadelain, Chimeric Antigen Receptor Therapy. N Engl J Med, 2018. 379(1): p. 64–73.

92. Denner, J. and R.R. Tonjes, Infection barriers to successful xenotransplantation focusing on porcine endogenous retroviruses. Clin Microbiol Rev, 2012. 25(2): p. 318–43.

93. Lu, T., et al., Xenotransplantation: Current Status in Preclinical Research. Front Immunol, 2019. 10: p. 3060.

94. Niu, D., et al., Inactivation of porcine endogenous retrovirus in pigs using CRISPR-Cas9. Science, 2017. 357(6357): p. 1303–1307.

95. Reardon, S., First pig-to-human heart transplant: what can scientists learn? Nature, 2022. 601(7893): p. 305–306.

96. Wilson, C.A., Porcine endogenous retroviruses and xenotransplantation. Cell Mol Life Sci, 2008. 65(21): p. 3399–412.

97. Suwanmanee, T., et al., Integration-deficient lentiviral vectors expressing codon-optimized R338L human FIX restore normal hemostasis in Hemophilia B mice. Mol Ther, 2014. 22(3): p. 567–574.

98. Cockrell, A.S. and T. Kafri, Gene delivery by lentivirus vectors. Mol Biotechnol, 2007. 36(3): p. 184–204.

99. Ma, H. and T. Kafri, A single-LTR HIV-1 vector optimized for functional genomics applications. Mol Ther, 2004. 10(1): p. 139–49.

100. Xu, K., et al., Generation of a stable cell line producing high-titer self-inactivating lentiviral vectors. Mol Ther, 2001. 3(1): p. 97–104.

